# Syntrophic microbiomes associated with methane-suppressive irrigation in rice

**DOI:** 10.64898/2026.06.15.732345

**Authors:** Kenny J.X. Lau, Ali Ma, Chen Bin, Thankaraj Salammal Maria Shibu, Srinivasan Ramachandran, Naweed I. Naqvi

## Abstract

Rice, a staple crop of nearly half of the world population, is grown predominantly in flooded paddies which are one of the largest contributors to methane emissions. An effective approach is to minimise the anaerobic flooded conditions that favour the growth of methanogenic archaea. Empirical measurements showed that controlled irrigation regime reduces methane emissions by 70% to 90%. The soil microbiomes of both flood and drip irrigated soil were characterised using whole-genome shotgun metagenomics. Controlled irrigation was shown to suppress methanogens and lower methane emissions. While emissions are correlated with *mcrA* gene abundance, empty flooded fields exhibited relatively high *mcrA* levels above baseline despite undetectable methane emissions. Rice cultivar genotype had no significant effect on the soil microbiomes. Co-occurrence network analysis indicates that soil microbial communities stratify according to their oxygen preferences along a gradient. Methanogens were increased in flooded paddies, and methane production attributed to the microorganisms involved in the anaerobic decay of organic matter. Controlled irrigation altered the microbiome by raising the soil redox potential by enhancing aeration and promoting ammonia oxidation and nitrification pathways.

**IMPORTANCE:** The temporal dynamics of microbial communities in drip-irrigated rice fields remain poorly characterized to-date. Empirical measurements demonstrate that controlled drip irrigation effectively suppresses methanogens and lowers methane emissions by up to 90%. Statistical analysis further revealed a moderate correlation between methane emissions and the *mcrA* gene with R = 0.6 and *p*-value = 2.9e-05. The correlation plot showed that the outliers corresponded to samples from empty flooded fields, where high *mcrA* gene abundance was observed despite low methane emissions. Methane produced in the soil is likely released into the atmosphere via transport through the aerenchyma of rice plants. Controlled irrigation is shown to be climate friendly as it reduces methane emissions by improving soil aeration and increasing the soil redox potential.

## INTRODUCTION

Traditional agricultural practices are being severely tested in the face of climate change. Rising temperatures, unpredictable precipitation patterns, and increasing occurrences of extreme weather events are prevailing challenges to global food security and will particularly impact the rice yields (1). Rice cultivation contributes to climate change through methane emissions from flooded paddies (2). Therefore, addressing these challenges require sustainable agricultural practices.

The choice of rice cultivars, irrigation methodologies and organic amendments such as rice straw impact methane emissions. Waterlogging in rice fields is known to increase methane emissions from rice fields (3). The anaerobic conditions in waterlogged paddies foster a conducive environment for methanogenic archaea to thrive therein, with methanogenesis being the final step in the anaerobic decay of organic matter. Providing more organic matter increases the substrates as carbon sources for methanogenesis (4). Secondly, rice cultivars with higher tiller numbers and longer roots provide more aerenchyma air spaces for the release of methane into the atmosphere (5). Root exudates such as fumarate are also thought to be contributors to methane emissions especially during the late stages of growth (6). Finally, water management is one of the most important factors where several studies have shown that draining flooded paddies in the mid-season or intermittently using alternate wetting and drying technique can effectively reduce methane emissions by 50% without incurring any yield losses (7).

Controlled irrigation can support sustainable rice cultivation by delivering water directly to the root zone of plants, thus reducing water wastage and ensuring optimal soil moisture levels (8). By incorporating drip irrigation into rice cultivation practices, farmers can enhance water use efficiency, reduce methane emissions, and contribute to a more sustainable agricultural system. It is well established that aerating the soil by draining out flooded paddies inhibits the growth of methanogens and hence reduces overall methane emission (9). However, there is limited knowledge regarding the changes in microbiota composition in the rhizosphere and the seasonal responses under aerobic controlled drip irrigation. The soil microbiome plays a crucial role in maintaining soil health and promoting plant growth and is responsible for nutrient cycling, decomposing organic matter, and even enhancing the resilience of crops to environmental stressors (10, 11). Plant physiology and their associated microbiomes may vary by genotype, three distinct semi-dwarf varieties, Temasek Rice (TR), Huanghuazhan (HHZ) and IR64 were selected to determine their comparative responses under controlled irrigation conditions.

TR was first developed in Singapore through marker-assisted breeding techniques, and it is high yielding, with dominant resistance genes against bacterial blight, blast, and drought/flooding stress (12). HHZ is a high-yielding rice variety developed in China and is known for its adaptability and resilience (13). It is widely cultivated in southern China and has been recognized for its tolerance to high temperatures and its ability to thrive in diverse environmental conditions. IR64 is a widely cultivated rice variety developed by the International Rice Research Institute (IRRI) in the Philippines and it is known for its high yield potential, early maturity, and resistance to diseases (14). Both HHZ and IR64 are used as baseline reference lines in rice breeding programs. Hence, in this study, these three indica semi-dwarf rice varieties were selected for comparison and whether there is a genotype-specific contribution to GHG emissions. We aimed to profile the methane levels and characterize the soil microbiomes of TR, HHZ, IR64 rice varieties and control plots without rice plants. The control plots are negative blank controls that provide information on the baseline soil microbiomes and the levels of methane emissions without rice plants. The trials were conducted in an outdoor rice field in Lim Chu Kang, Singapore. Rhizosphere soil samples were collected at one-month intervals during the early (week 4), mid (week 8), and late (week 12) post-transplantation growth stages of rice for metagenomic sequencing.

## MATERIALS AND METHODS

### Soil preparation and methane measurements

Soil was prepared by mixing topsoil with 30% (v/v) peat moss (BVB, Netherlands), sheep manure at 300 g/m², and chopped rice straw at 1000 g/m², which were then blended thoroughly. The mixed soil was then added into outdoor plots of 1.5 m × 1.5 m. Temasek Rice, HHZ and IR64 rice seeds were sown in the greenhouse. Twenty-day old rice seedlings were then transplanted into the field plots at an outdoor site at Lim Chu Kang, Singapore (103⁰70’49’’ E and 1⁰42’73’’ N). The three rice varieties were subjected to drip and flood irrigation. Drip irrigation plots had six lateral pipelines to provide water and fertilizer at frequency of two minutes per cycles. Six plants were planted evenly spaced out on each lateral drip line with a total of 36 plants grown in each plot. The continuous flooded plots had rice plants that were submerged in flood water to a depth of 10 cm above the soil level.

The static closed chamber method was used to measure methane (CH_4_) by placing a three-part chamber, the bottom chamber base installed over the soil. The middle chamber is added when rice plants mature and become taller. An electric fan was fixed in the top compartment of each acrylic transparent chamber of 50 cm × 50 cm × 50 cm with a rubber sampling window at the side of the chamber to facilitate the withdrawal of air sample through a needle and syringe. 20 mL of air was drawn using a 25 mL syringe at 0, 10, 20 and 30 minutes. Methane sampling was conducted biweekly between 1000h to 1200h. Methane measurements were then analysed using a gas chromatograph with a flame ionization detector, GC-2010 Pro (Shimadzu, Japan). The methane flux was determined by the following formula:

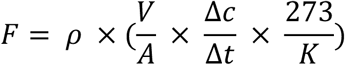

F is CH_4_ flux expressed in (mg CH_4_ m^-2^ h^-1^), ρ is the gas density of CH_4_ gas (0.174 mg cm^-3^), V is the volume of the chamber in (m^-3^), A is the surface area of the chamber in (m^-2^), 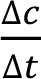 is the rate of gas concentration increase in the chamber (mg m^-3^ day^-1^) and K is air temperature in the chamber measured in Kelvin. Gas chromatography-flame ionization detector (GC-FID) was performed using the Poraplot Q-HT column with carrier helium gas flow rate of 18 ml/min, column temperature of 90 ⁰C, FID at 150 ⁰C, hydrogen at 40 ml/min and air at 400 ml/min. Readings obtained were then converted to mg/m^2^ day and plotted as boxplot using *ggplot2* (15) in R.

### DNA extraction, library preparation, sequencing and bioinformatics

Soil samples were collected by uprooting the plants and collected the root soil in 15 ml phosphate saline buffer (PBS) in a 50 ml tube. The 50 ml tube with soil and roots were sonicated at 50-60Hz for 5 minutes. The tube was then centrifuged at 4000 Xg for 1 minute and supernatant was discarded leaving behind the soil. 1 g of soil was weighed and extracted using the DNeasy Power Soil Pro kit (Qiagen, Germany) following manufacturer’s protocol. 100 µL of extracted DNA was then stored at -20 ⁰C freezer for next generation sequencing. Quantitative polymerase chain reaction (qPCR) was performed using primers targeting the *mcrA* gene using the Met630F GGATTAGATACCCSGGTAGT and Met803R GTTGARTCCAATTAAACCGCA primers. The qPCR mixture consisted of 1 µL of DNA template and 0.25 µM of primers with 2 × master mix. Negative controls were tested using sterile water. qPCR was performed using the Kapa Sybr Fast Universal qPCR kit (Roche, USA) in CFX96 thermal cycler (BioRad, USA) with 40 cycles of two-step amplification at 95 ⁰C for 10 seconds and 60 ⁰C for 40 seconds followed by a melting curve. Standard curve for qPCR analysis was quantitated by performing serial dilution of a gBlock containing with known 10^10^ copies of the *mcrA* gene of *Methanobacterium* as follows: CCA GGG GCG CGA ACC GGA TTA GAT ACC CGG GTA GTC CTG GCC GTA AAC GAT GCA GAC TTG GTG TTG GGA TGG CTT CGA GCT GCT CCA GTG CCG AAG GGA AGC TGT TAA GTC TGC CGC CTG GGA AGT ACG GTC GCA AGA CTG AAA CTT AAA GGA ATT GGC GGG GGA GCA CCA CAA CGC GTG GAG CCT GCG GTT TAA TTG GAT TCA ACG CCG GAC synthesized by Integrated DNA Technologies (IDT, USA). Gene copy number abundance was determined based on the straight-line equation of Cq versus log of known copies of the standard curve template. The gene copy numbers were normalised to 1 g of soil.

High throughput sequencing libraries were prepared using the TruSeq Nano DNA kit (Illumina, USA). Whole genome shotgun metagenomic sequencing was performed at Macrogen, Singapore, using the NovaSeq 6000 sequencer. Sequence adapters were trimmed using Trimmomatic (16) paired end reads using Illumina Clip step and TruSeq3 adapter sequences with a phred score threshold of 20 (**Table S1**). Reads were then queried across the NCBI-nr database (28 July 2023) using Diamond v2.1.10 (17) running in BLASTx mode with the default BLOSUM62 substitution matrix. The resulting outputs were DAA formatted files and were parsed into Metagenome Analyzer v6.24.22 (18) for analysis. Taxonomic classification was performed at genus level, and the taxonomic tables were exported as txt files for downstream analysis using *vegan* (19) and *phyloseq* (20) packages in R. Alpha diversity was assessed using Shannon (H) and Simpson (D) diversity indices. Principal coordinate analysis (PCoA) was then plotted to identify clustering patterns with metadata and environmental factors. Differential taxa were identified using *MaAsLin2* (21) package with irrigation and variety defined as fixed factors using the default parameters.

Taxonomic composition at the treatment level was evaluated by aggregating replicates by flood and drip treatment group using merge_samples function in R. Raw count values were then transformed into relative abundances by dividing the count of each taxon by the total number of reads within the flood and drip irrigation group. A bipartite co-occurrence network was constructed using the R package igraph (22). Nodes represented individual taxa while edges denote the presence and relative abundance of a taxa within the flood or drip group. Edge-weight filtration step was performed to remove edges corresponding to a taxa relative abundance of less than 0.5%. Nodes that have a degree of 0 with no neighbours were also removed from the graph. Nodes were then annotated and coloured based on the first-degree neighbour connections of flood or drip neighbours. The annotated network was exported into Cytoscape 3.10.4 (23) via the RCy3 package (24) for visual rendering.

Lastly, the reads were assembled into Metagenomes-Assembled Genomes (MAGs) in KBase with MegaHIT (25) and Concoct v1.1 (26) using a minimum contig length of 2500 bp. Bins were extracted as assemblies and filtered using CheckM (27) and bins were taxonomically classified with GTDB-Tk v2.3.2 (28). The MAGs were then functionally annotated using DRAM (29) in KBase.

### Soil redox potential measurements

The pH, ORP and temperature meter probe of HI991003 (Hanna Instruments, Italy) was inserted to measure the soil redox potential. The probe was left in the soil for 5 minutes until the readings were stabilized before recording.

## RESULTS

### Correlating Methane Emissions with *mcrA* Gene Copy Numbers

Methane emissions were empirically measured from flood- and drip-irrigated field plots using the static closed chamber method coupled with gas chromatography. Based on these results, we observed that there is a significant reduction of 70-90% in methane emissions when drip irrigation was applied to rice plants under field conditions (**Fig. 1, Table S2**). Field measurements were conducted in triplicates and reduction in methane was consistently observed across three different rice varieties under the aforementioned drip irrigation. Two emission peaks were identified at Week 3 and Week 9, which coincides with tillering and flowering stages of plant development. Additionally, Huanghuazhan appears to have a slightly lower methane emission than IR64 and Temasek rice. A small peak was observed at week 5 in the flooded control plot without plants, compared to the drip-irrigated control plot without plants. Baseline methane emissions in control plots without rice plants ranged from 0.6 to 57.8 mg/m² per day. A moderate correlation of R = 0.6 and *p*-value = 2.9e-05 was observed between methane emission and *mcrA* gene copies per gram dry soil (**Fig. 2**). Our result suggests an exponential increase in the levels of methanogens with the highest *mcrA* abundance observed in flooded fields after 3 months post transplantation (**Table S3**). *mcrA* gene copy number remains relatively low with slight increase in abundance in drip irrigated soils towards the end of the season. Notably, the control plots without rice plants emit relatively low levels of methane but the *mcrA* gene abundance remains equally as high as those in the soil grown with rice plants.

**Figure 1.**
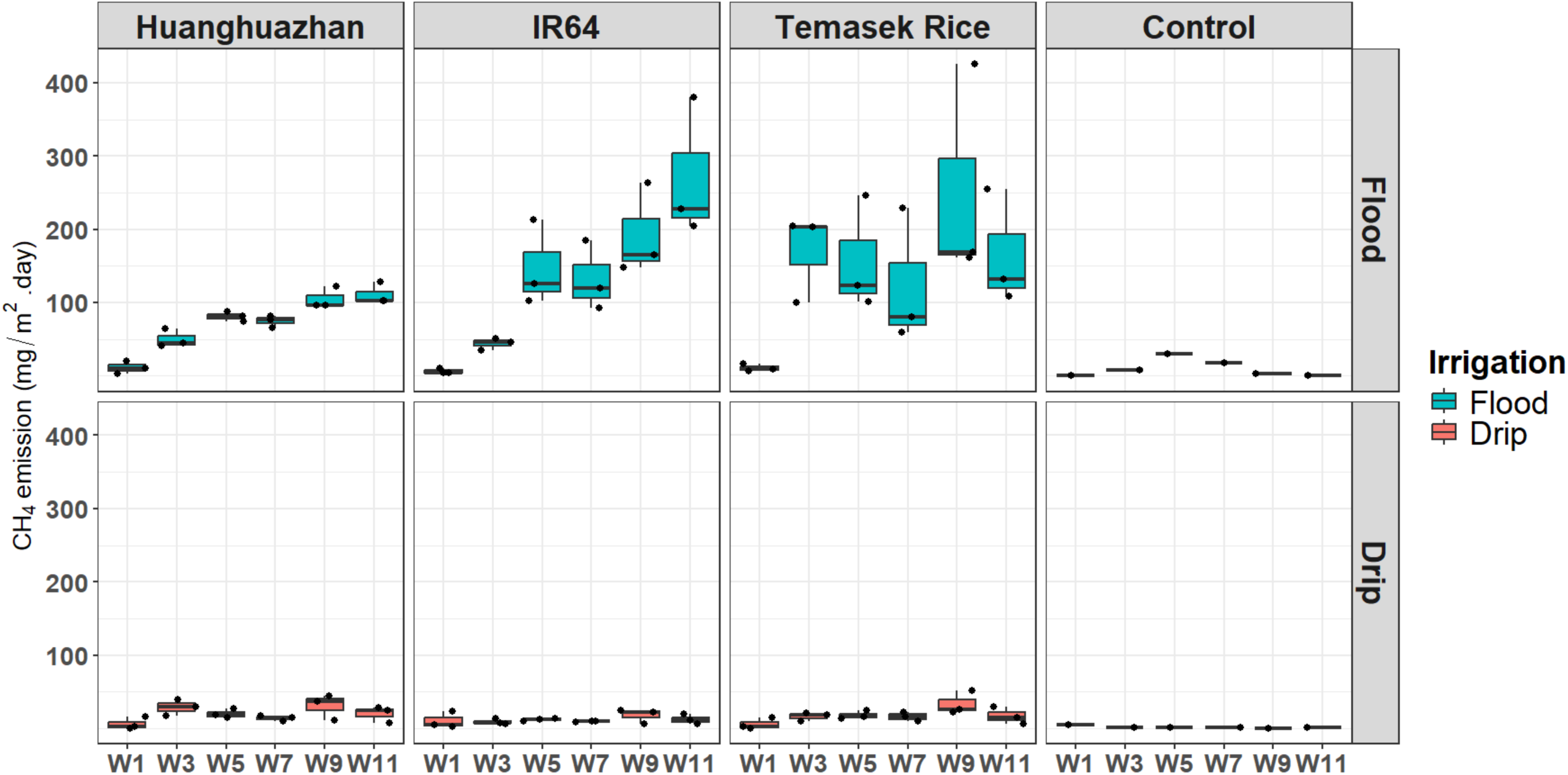
Comparison of methane emissions from three rice varieties (Temasek Rice, Huanghuazhan, IR64) and control plots under flood versus drip irrigation. The top panel illustrates emissions under flood irrigation, while the bottom panel shows emissions under drip irrigation. A significant reduction of approximately 70-90% in methane emissions was observed for all rice varieties with drip irrigation with triplicates. Blank control plots represent baseline methane emissions from flooded or drip-irrigated soils without rice plants.

**Figure 2.**
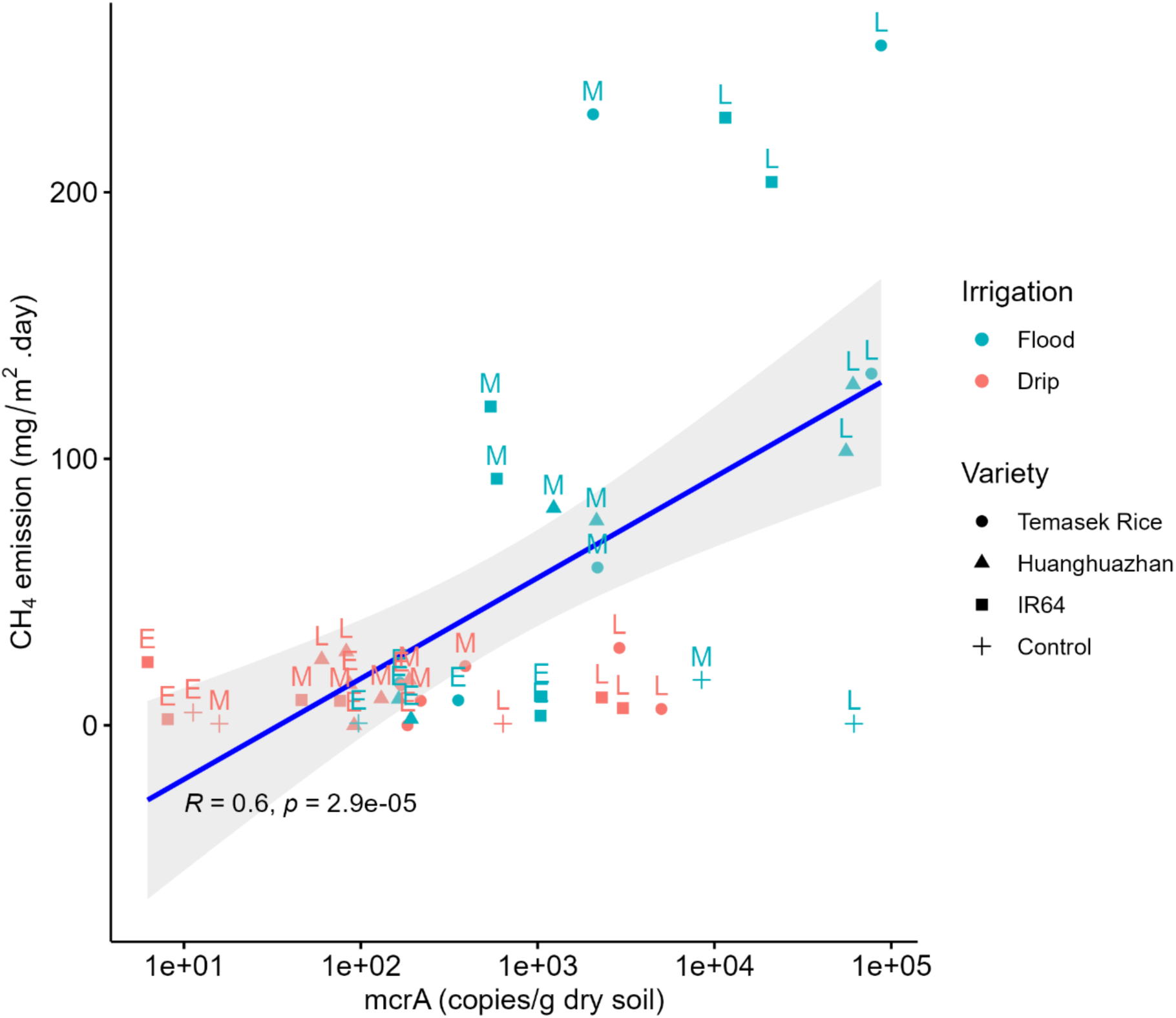
Correlation plot of methane emission versus *mcrA* gene copy number estimates. Methane emissions and its corresponding *mcrA* gene copy number from soil samples collected at Early (E), Mid (M) and Late (L) season timepoints. A moderate correlation was observed with a Pearson’s R of 0.6 and p-value of 2.9e-05, where correlation between levels of methane emissions and *mcrA* gene abundance in continuous flooded soils are higher than drip-irrigated soils. In contrast, control plots without plants have relatively low methane emissions despite having relatively equal *mcrA* gene abundance levels as the plots with rice plants.

A total of 24 samples were collected for metagenomic sequencing. The sequencing run yielded an average of 81,323,058 reads for metagenomic analysis. Sequencing reads and binning information for this study can be found in the supplemental data and dryad depository at https://doi.org/10.5061/dryad.vt4b8gv67. Samples were collected at 1-, 2-, and 3-months post-transplantation to monitor longitudinal changes in the microbiome over time. Based on the Shannon and Simpson indices in **Table S4** suggests that HHZ has a slightly lower soil microbial diversity in the early and mid-stage samples. Soil has increasing microbial diversity towards the late time point and peak at the highest after 3 months post-transplantation. Control plots without rice plants clustered alongside both flood- and drip-irrigated rice samples (**Fig. 3**). The results indicate that anaerobic degradation of organic matter plays a dominant role in shaping the soil microbiome, with rice plant exerting a comparatively minor influence within the rhizosphere under the conditions of this study. The relative abundance of methanogens increased gradually, where they peak at around three months post-transplantation during the rice ripening stage (**Fig. 4, Fig. S1**). *Methanosarcina* was detected in both flood and drip samples but exhibited higher concentrations under flood-irrigated conditions. The other three methanogenic genera, *Methanobacterium*, *Methanocella*, and *Methanothrix* were higher in abundance at the late time points during maturation as compared to the early time points and they play crucial roles in the global carbon cycle through the production of methane.

**Figure 3.**
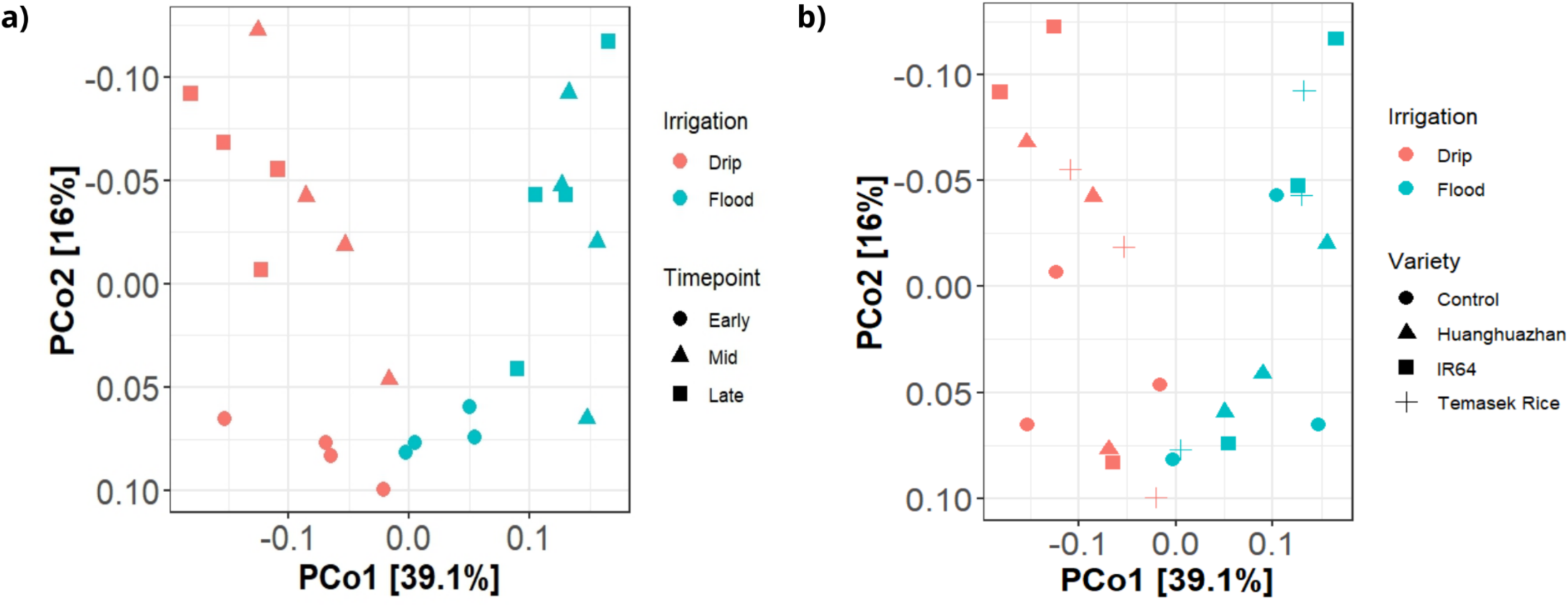
Clustering analysis of soil rhizosphere microbiomes over the growing season. The microbiomes diverge into distinct clusters, primarily along the PCo1 axis, as the plant matures from early to late season. **a)** Samples are coloured by irrigation type and shaped by rice variety. **b)** Temporal clustering is evident, with sample shapes representing collection dates. Separation along the PCo1 axis distinguishes drip-irrigated from flooded soils, with early-season samples exhibiting greater similarity compared to those from mid- and late-season. Notably, rice plots and control plots without rice plants show similar clustering patterns, indicating no significant difference attributable to the presence of rice plants.

**Figure 4.**
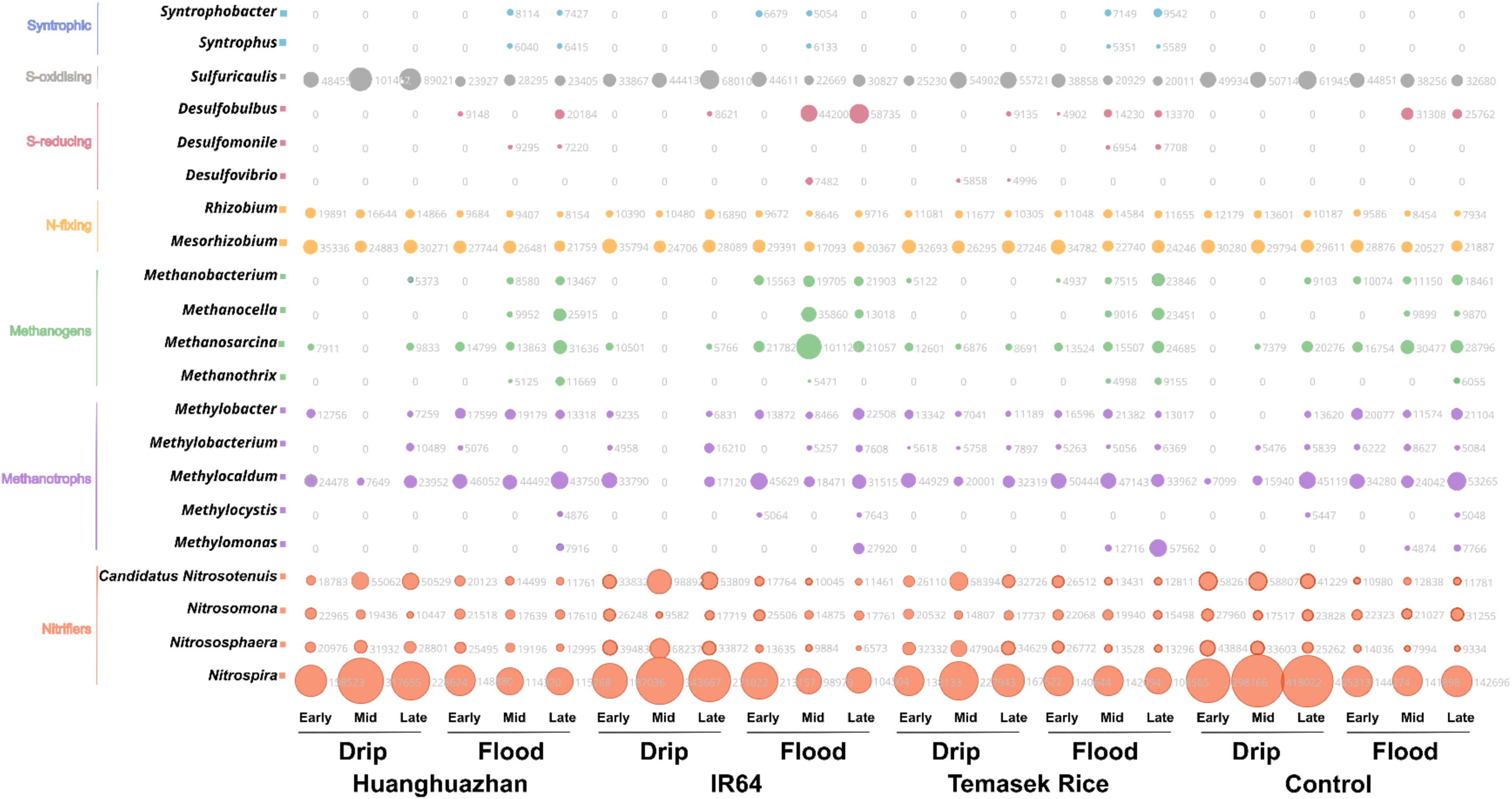
Shifts in the relative abundance of nitrifiers, methanotrophs, methanogens, sulphur-oxidising and sulphur-reducing taxa were observed in the soil microbiomes under drip and flood irrigation for Huanghuazhan, IR64, Temasek Rice and Control plots. Drip irrigation was associated with higher relative abundance levels of nitrifiers and sulphur oxidisers, while flood irrigation promoted the proliferation of methanogens and sulphur-reducing taxa.

### Relationship between Methane Flux and Differentially Abundant Microbial taxa

Correlation analysis was conducted between methane emissions and microbial taxa exhibiting statistically significant differential abundance in drip-versus flood-irrigated soils. The top panel of **Fig. 5** highlights four methanogenic genera, *Methanobacterium*, *Methanocella*, *Methanothrix*, and *Methanosarcina*, that were positively correlated with methane emissions. Stronger correlations were observed for hydrogenotrophic methanogens (*Methanobacterium*: R = 0.62; *Methanocella*: R = 0.59), whereas acetoclastic methanogens (*Methanothrix*: R = 0.4; *Methanosarcina*: R = 0.37) showed weaker associations. *Methanothrix*, a specialist acetate-utilizing methanogen, likely showed reduced correlation. In contrast, *Methanosarcina*, a metabolic generalist capable of hydrogenotrophic, acetoclastic, and methylotrophic methanogenesis, is the most abundant methanogen, but weak correlation was found due to their generalist behaviour and outlier data point observed at week 7.

**Figure 5.**
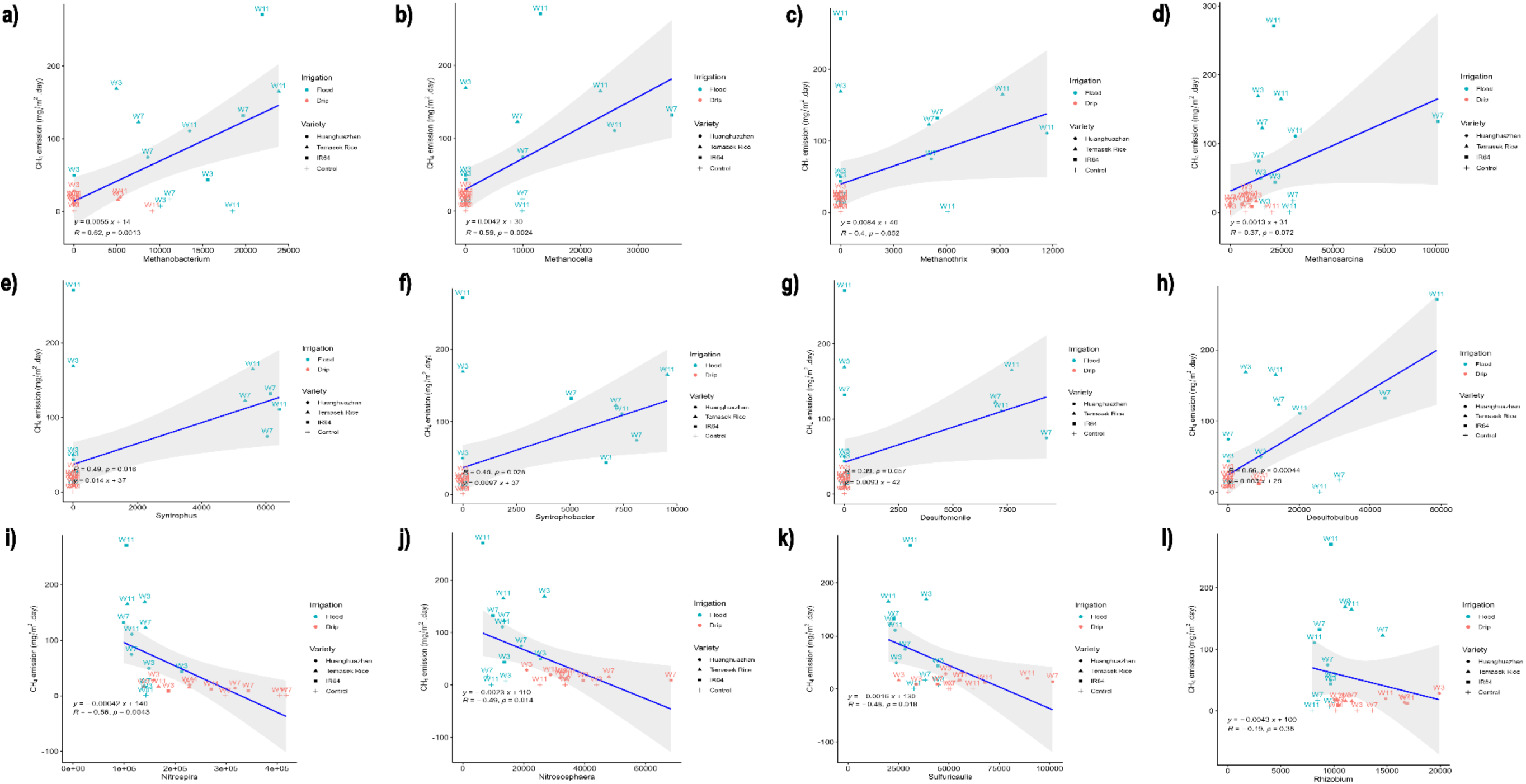
Correlation analysis between methane emissions and microbial taxa. Top panel **(a-d)** are methanogens namely *Methanobacterium*, *Methanocella*, *Methanocella* and *Methanosarcina*, middle panel **(e)** and **(f)** are syntrophic bacteria, middle panel **(g)** and **(h)** are sulfur-reducers that show positive correlations with methane emission. Conversely, bottom panel **(i)** and **(j)** are nitrifiers, **(k)** sulfur-oxidiser, and **(l)** nitrogen fixer exhibit negative correlations with methane emissions.

Additionally, syntrophic bacteria such as *Syntrophus* and *Syntrophobacter*, along with sulphate-reducing taxa including *Desulfomonile* and *Desulfobulbus*, were also positively correlated with methane emissions. These methanogenic, syntrophic, and sulphur-reducing taxa were enriched under continuously flooded conditions. Conversely, *Nitrospira*, *Nitrososphaera*, *Sulfuricaulis*, and *Rhizobium* were negatively correlated with methane emissions, indicating higher relative abundance in drip-irrigated soils. These taxa representing nitrifiers, sulfur oxidizers, and nitrogen fixers were more prevalent under aerobic conditions, supporting the hypothesis that drip irrigation promotes nitrification and sulfur-oxidising related members in the microbial communities adapted to aerobic environments. *Nitrospira* and *Sulfuricaulis* were observed to be higher in the drip-irrigated plots between mid and late stages of rice development.

### Irrigation regimes shape microbial communities along an oxygen gradient

Increased soil oxygen levels in drip irrigation have favoured the proliferation of *Nitrospira* and *Sulfuricaulis*, in contrast to flooded conditions that support the dominance of methanogens, syntrophic and sulfur-reducing microbes. FAPROTAX functional gene predictions also suggest that flood-irrigated soils are typically enriched in microbes performing methanogenesis, methanotrophy and methylotrophy-related activities (**Fig. S2**). This is consistent with anaerobic conditions found in flooded soils promoting methanogens. In drip-irrigated soil, methane-related functions are reduced significantly. Nitrogen-cycling groups become proportionally larger where the nitrification and aerobic ammonia oxidation are enriched, suggesting that aerobic conditions favoured these pathways. Temporal shifts in functional pathways from early to late time points are also evident as the relative proportions of FAPROTAX functions change and become more prominent at the late time points. Soil redox potential measurements collected in the fields were in the typical range of 0 to 150 mV in drip-irrigated and -50 to -200 mV in flood-irrigated soils (**Fig. 6a**). Positive ORP range suggests oxidized state favouring nitrifiers and ammonia oxidation-related activities while negative ORP range suggests anaerobic state, favouring methanogenesis processes.

**Figure 6.**
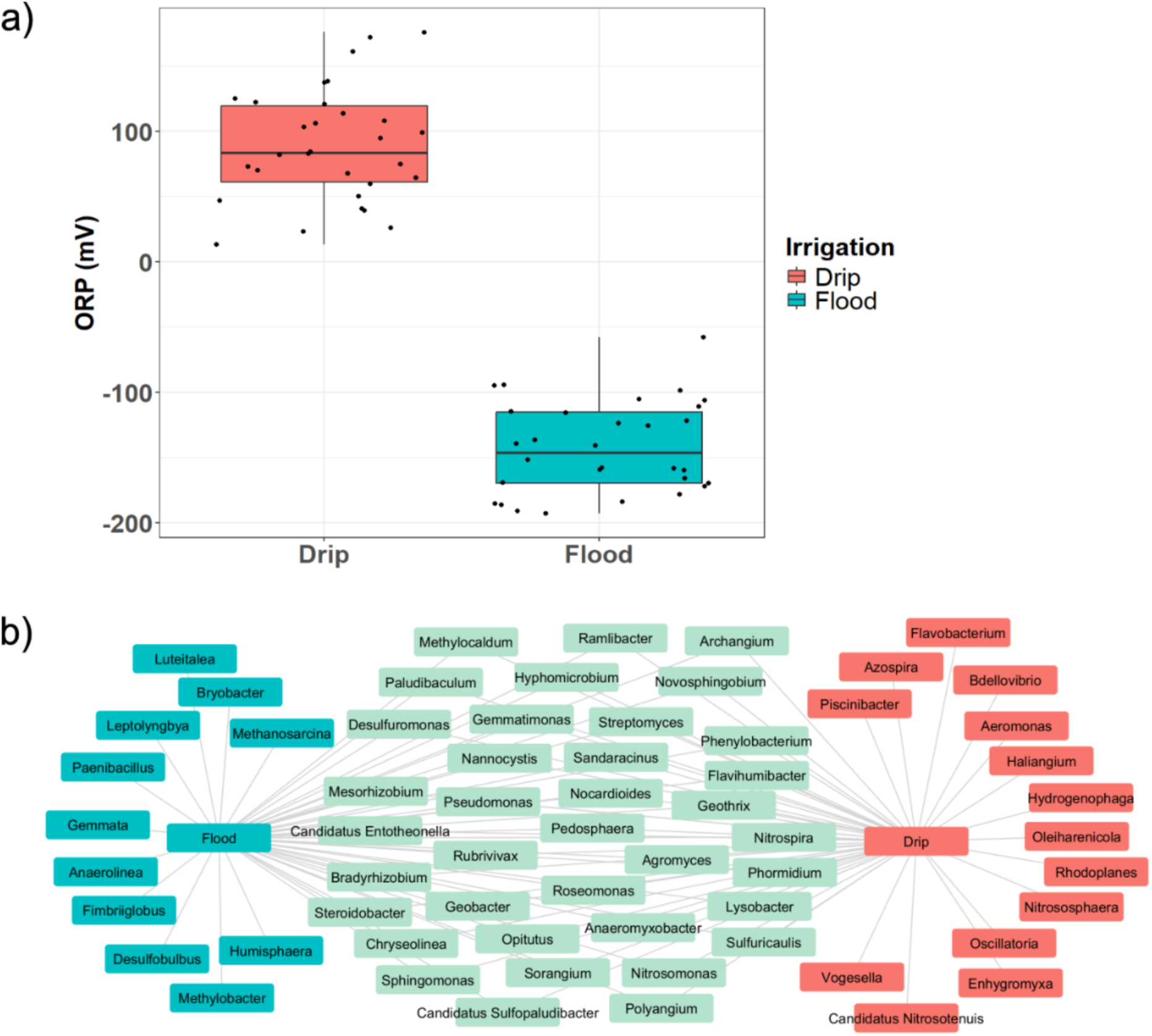
Oxygen availability influences the soil redox potential and selects for differential taxa. **a)** Drip irrigation yields a higher soil redox potential compared to continuous flooding. Box plot showing ORP readings of drip and flood-irrigated soils measured over 12 weeks with triplicate samples collected biweekly. **b)** Co-occurrence networks reveal microbes enriched under flood and drip irrigation. Flooded conditions favour the proliferation of *Methanosarcina*, *Methylobacter*, *Anaerolinea*, *Humisphaera*, *Paenibacillus*, *Desulfobulbus* and *Gemmata*. Drip irrigation enriches primarily for aerobes and facultative aerobes including *Nitrosophaera*, *Oscillatoria*, *Aeromonas*, *Candidatus Nitrosotenuis*, *Vogsella* and *Azospira*. Several taxa are shared between both irrigation regimes, and the network reveals a gradient from flood-associated microbes to drip-associated microbes.

Co-occurrence networks revealed distinct microbial clusters associated with flood and drip irrigation regimes. **Fig. 6b** shows that under flooded condition, the networks shift towards a dominance of anaerobic and methanogenic bacteria with increased proliferation of *Methanosarcina*, *Methylobacter*, *Anaerolinea*, *Humisphaera*, *Paenibacillus*, *Desulfobulbus* and *Gemmata*. The oxygen-limited environment favours the obligate and facultative anaerobes. For example, methanogens and sulfur-reducers do not require oxygen as electron acceptors and *Paenibacillus* can form endospores to survive in oxygen-poor soils. In contrast, drip irrigation was characterized by the enrichment of aerobic and facultative anaerobic taxa, including *Nitrosophaera*, *Oscillatoria*, *Aeromonas*, *Candidatus Nitrosotenuis*, *Vogesella* and *Azospira*. Many taxa are shared between both irrigation regimes and they are considered generalists. Their survival in both flood and drip environments can be attributed to their high degree of ecological plasticity across varying soil moisture and their metabolic versatility in switching between aerobic and anaerobic states. For instance, *Methylocaldum* and *Ramlibacter* can form cysts to allow them to endure prolonged in oxygen-depleted environments and desiccation. The network topology revealed a microbial preferential gradient where flood-associated taxa tend to cluster towards the left of the network and drip-associated taxa positioned towards the right, suggesting a transition in the redox and oxygenated state of the soil.

## DISCUSSION

### Irrigation regime reshapes methanogenic and syntrophic networks

Among the methanogens, *Methanosarcina* was found in higher abundance in the flood-irrigated soil and is considered a generalist microbe as it can utilise multiple methanogenic biosynthesis pathways (**Fig. 4**). *Methanosarcina* thrives at higher temperatures and may potentially outcompete strict acetoclastic *Methanothrix* if acetate becomes depleted (30). *Methanothrix* specializes in converting acetate (CH₃COO⁻) into methane and carbon dioxide and is particularly significant in many anaerobic ecosystems due to its high affinity for acetate (31). *Methanobacterium* species was also detected in flooded soil. This is a well-characterized hydrogenotrophic methanogen, which primarily utilizes hydrogen (H₂) as an electron donor to reduce carbon dioxide (CO₂) to methane (CH₄) and is commonly found in diverse anaerobic environments like sediments, wastewater, and animal digestive tracts [32]. Flood-irrigated soil also harbours hydrogenotrophic methanogens, *Methanocella*, formerly Rice Cluster I (**Fig. S1**). *Methanocella* is classified as hydrogenotrophic methanogen that also relies on the H₂/CO₂ pathway and is particularly noted for prevalence in environments like rice paddies (33). Additionally, *Syntrophobacter* and *Syntrophus* were enriched in flooded plots, contributing to acetate precursor production from propionate, thereby supporting methanogenic growth (34). *Syntrophobacter* and *Syntrophus* are genera of syntrophic bacteria that are essential for the anaerobic breakdown of complex organic matter. They are not methanogens themselves but obligate partners that perform the initial steps of anaerobic degradation. *Syntrophobacter* species is particularly known for the ability to oxidize propionate into acetate, CO₂, and hydrogen or formate. This conversion is thermodynamically favourable when hydrogen and formate concentrations in soil are low and has been demonstrated with ^13^C stable isotope experiments previously (35). Additionally, *Syntrophobacter* was found to be closely associated with methanogens and are the most dominant propionate-oxidising syntrophic bacteria in rice fields (36). Similarly, *Syntrophus* species perform syntrophic degradation of other fatty acids like butyrate and aromatic compounds such as benzoate, also producing acetate, CO₂, and hydrogen and formate that are subsequently consumed by their metabolic partners (37). Both genera are therefore key intermediary anaerobes, distributing carbon from more complex organic molecules towards substrates usable by acetoclastic and hydrogenotrophic methanogens, eventually leading to methane formation in environments like anoxic soils. Studies have examined the dynamics of methane production alongside shifts in methanogenic communities during rice growth (38, 39). They show that during initial stages of rice cultivation, methane emissions are typically low. As the plants develop and anaerobic conditions persist, methane fluxes tend to increase progressively over time (40). On the other hand, drip-irrigated soils harboured a higher abundance of nitrifier species compared to flooded soils, with *Nitrospira* emerging as the dominant nitrifying taxon under drip irrigation. The relative abundance levels of methanogens and *Nitrospira* vary due to oxygen availability in the soil. The process is primarily driven by the change in redox potential status of the soil (41). As seen previously in **Fig. 6a**, negative ORP range implicates a shift in the microbiomes towards methanogenesis-dominant functional pathways in anaerobic flooded condition while positive ORP range shifts towards nitrification-dominant pathways under drip conditions where microbial populations of methanogens decrease and *Nitrospira* population increase over time.

Statistical analysis using *MaAsLin2* showed a list of identified taxa that were significantly differentiated in the irrigation groups (top 16 listed in **Table S5**, based on lowest *p*-values). *Usitatibacter*, *Candidatus Nitrosotenuis*, *Nitrospira*, and *Pseudomonas* were significantly enriched in drip-irrigated soils. Conversely, *Aquisphaera*, *Gemmata*, *Caldilinea*, and *Methanosarcina* were more prevalent in flood-irrigated soils. These distinct microbial enrichments likely reflect differences in environmental filtering pressures, particularly the oxygen-limited conditions characteristic of flooded systems. Flooded paddies also exhibit an enrichment of sulfur-reducing bacteria, including *Desulfobulbus*, *Desulforomonas*, and *Desulfomonile*. *Usitatibacter* has been identified as a key species after long-term manure and straw amendments and is reported to help mediate soil carbon and phosphorus cycling in a separate study (42). *Candidatus Nitrosotenuis* is known to convert ammonia to nitrite (43) while *Nitrospira* converts nitrite to nitrate and both microbes play an important role in the Nitrogen cycle (44). *Pseudomonas* genus encompasses a wide range of bacteria, including *P. denitrificans* which functions as a de-nitrifier by reducing nitrate to gaseous nitrogen compounds under anaerobic conditions. In contrast, the other *Pseudomonas* species are primarily involved in the decomposition of organic matter, releasing nutrients like nitrogen, phosphorus and sulfur back into the soil (45, 46). *Aquisphaera*, *Gemmata* and *Caldilinea* are reported to play a role in degrading organic matter and cycling of nutrients in the soil (47–49). *Methanosarcina* is one of the key methanogenic archaea that contributes to methane emissions. *Methanobacterium* primarily reduces carbon dioxide with hydrogen to produce methane and *Methanothrix* specializes in acetoclastic methanogenesis. Unlike *Methanobacterium* and *Methanothrix*, *Methanosarcina* can utilize multiple substrates and is capable of both acetoclastic and methylotrophic pathways (30). Methanogens associate themselves close to the root and methane is likely channelled through the aerenchyma cells into the atmosphere.

### Reduced Tillering and Efficient Grain Carbon Partitioning as Low-Methane Traits

Based on our results in **Fig. 1**, field plots cultivated with rice plants emitted about 20 to 50 times higher methane than the control empty plots without plants. It is thought that the root aerenchyma cells provide a route for methane to escape from the waterlogged soil. Positive correlation between tiller number and methane release was found in a study suggesting that each tiller could function as a conduit for methane transport in flooded paddies (50). Longitudinal monitoring of methane flux across key developmental stages of rice, booting, heading, grain filling, and maturity also revealed an exponential increase in methane emissions during the heading stage (51). Comparative study among varieties such as B40, IR64, and IR72 further support these observations, where IR64 and IR72 produce more tillers and consequently emit higher methane levels than B40 (52). Besides reduced tillers, previous studies indicate that the harvest index, defined as the ratio of grain yield to total above-ground dry matter, is also critical for coupling high yields with low methane emissions (53, 54). A high harvest index demonstrates that carbon is preferentially allocated to the grain sink rather than the root system. HHZ, a semi-dwarf cultivar, is characterized by a slightly taller plant height, more grains per panicle, and fewer tillers has a higher harvest index. In contrast, Temasek Rice and IR64, also semi-dwarf varieties, exhibit shorter plant height, a lower seed setting rate, and slightly higher tiller density. Consistent with these findings, our data in **Fig. 1** showed that HHZ emitted slightly lower methane levels than Temasek rice and IR64. This is likely attributable to HHZ having reduced tillers and higher seed settling rate relative to Temasek Rice and IR64 varieties (**Table S6**). This observation further supports the hypothesis that methane emissions are regulated by two key plant traits, namely the transport capacity of the aerenchyma and the efficiency of carbon allocation to the grains. Therefore, low emission rice cultivars should typically have traits such as reduced tillers, smaller aerenchyma to stele ratio and high harvest index.

### Draft genomes reconstructed from metagenomic data reveal distinct functional taxonomic bins in flood- and drip-irrigated soils

The co-occurrence clusters revealed only potential pairwise associations based on Spearman correlations. To further assess whether complete microbial genomes were represented, metagenome-assembled genomes (MAGs) were reconstructed from soil samples. Taxonomic classification using GTDB-Tk resolved most bins down to the family level, while half of the assemblage remained unclassified. Notably, methanogens and methanotrophs were not recovered in any of the MAGs bins, likely because their read coverage were too low to assemble into longer contigs. Both flood and drip-irrigated soils share some similar MAGs that include *Anaerolineales*, *Desulfuromonadales*, *Ignavibacteriaceae*, *Deferrimicrobiaceae* and *Steroidobacteriaceae* (**Fig. S3** and **Fig. S4**). *Anaerolineales* was reported to cooperate syntrophically with *Methanosaeta* in the degradation of alkanes in soil (55). In this process, *Anaerolineales* activate alkanes through fumarate addition and degrade them into fatty acids and acetate (56) and *Methanosaeta* then use the acetate to synthesize methane and carbon dioxide (57). *Ignavibacteriaceae* are facultative anaerobes and they degrade organic matter and perform anaerobic respiration with nitrate as electron acceptor (58). This observation also tallies with previous work that showed its association with *Geobacter* and *Anaerolineae* in methane production clusters (59). **Fig. S3** shows MAGs associated with flood irrigation and enrichment for *Phormidium* (bin.035), *Woesarchaeales* (bin.028), *Bacteroidales* (bin.021) and *Verrucomicrobiaes* (bin.008). *Phormidium* are photosynthetic cyanobacteria and are found in relatively large numbers in paddy flood water. *Phormidium* are beneficial for mature rice plants where they fix nitrogen and release growth-promoting phytohormones such as auxins and gibberellins to enhance root growth (60). *Woesarchaeales* are ubiquitous in anoxic environments such as oil rigs, wastewater systems, paddy soils and sulfuric springs and are thought to be involved in anaerobic carbon-nitrogen-sulphur cycling, likely playing a role as fermentative syntrophs that exchange metabolites with other microbial partners (61). MAGs belonging to *Myxococcota* (bin.034 and 048), *Methylomirabilales* (bin.039) and *Nitrosotenuis* (bin.046) were recovered from the drip-irrigated soil (**Fig. S4**). *Myxococcota* have been reported in mangrove soil, and they probably thrive under moist and suboxic environments like the drip-irrigated soil (62). *Methylomirabilales* MAGs were reported to be found in oxic soil depths with methylotrophy functions, and they do not have methane monooxygenase or nitrogen reduction genes (63). Likewise, we also identified *Methylomirabilales* in the drip-irrigated soils. *Methylomirabilales* can perform nitrite-dependent anaerobic methane oxidation (n-damo) in low oxygen environments (64). This process may be supported by *Nitrosotenuis*, an ammonia-oxidising archaeon that converts ammonia into nitrite required for nitrite-dependent anaerobic methane oxidation (N-DAMO) (43). *Nitrosotenuis* are higher in abundance in drip-irrigated soil. Additionally, *Burkholderiales* and *Rhodocyclaceae* can grow in both aerobic and anoxic conditions. Under anaerobic conditions, they can function as denitrifiers and might compete with *Methylomirabilales* for nitrite. Despite the recovery of MAGs from both flood and drip irrigation regimes, a significant portion of the metagenome remains unclassified. To achieve a systematic understanding of soil microbial dynamics, future research will aim to integrate microbial, biogeochemical and greenhouse gas modelling with time-resolved transcriptomic analyses to gain mechanistic insight into the highly complex syntrophic interactions associated with and leading up to microbial mitigation of climate impact.

## CONCLUSIONS

Harnessing soil microbiome knowledge within sustainable rice cultivation offers a dual benefit. It can improve crop productivity by enhancing nutrient cycling and plant resilience, while simultaneously mitigating greenhouse gas emissions and reducing environmental impact. Empirical findings of the microbiomes of plots with and without rice plants suggest that methane production is primarily a result of anaerobic decomposition of organic matter as the soil microbiomes remain relatively similar even in the absence of rice plants. Our findings add to prior reports indicating that approximately 90% of methane produced is released through plants. Since methane is insoluble in water, it is likely trapped in the soil and is released through openings like the aerenchyma and rice tillers into the atmosphere. This accounts for the observed peaks in methane emission profiles during the tillering, heading, and ripening stages of rice growth, when tiller numbers increase and undergo changes. Controlled irrigation thus enhances soil aeration, reshapes the rhizosphere microbiomes and consequently reduces methane emissions by a significant and impactful quantum.

## DATA AVAILABILITY

Raw metagenomic sequencing data have been submitted to NCBI under BioProject ID of <u>PRJNA1377271</u>. All 24 metagenome sequence files used in the analyses are publicly available, with accession numbers ranging from SRX31415382 to SRX31415405.

Metagenome assembled contigs and taxonomic bins data are deposited in a Dryad repository at https://doi.org/10.5061/dryad.vt4b8gv67

Metagenome-Assembled Genomes bioinformatics analyses are available at this KBase narrative at https://narrative.kbase.us/narrative/236707

## ACKNOWLEDGEMENTS

We thank Zhong Chao Yin for sharing the rice varieties and providing helpful suggestions throughout the project. We acknowledge project management support from Phuay Yee Goh. We thank the Macrogen team for metagenomic sequencing. This research was funded by grants from the Philanthropy Asia Alliance, the Gates Foundation, and the Temasek Life Sciences Laboratory, Singapore.

## AUTHOR CONTRIBUTIONS

NIN, SR and KJXL designed the experiments and microbiome analysis framework. KJXL performed the soil collection, root imaging, DNA extraction, microbiome data analysis, bioinformatics and statistical analyses. Methane emissions analysis was performed by AM, CB and KS. AM and TSMS cultivated and maintained the rice plants. NIN supervised KJXL on both the research work, experimental design, data analyses and manuscript writing. NIN and KJXL analysed the microbiome data and co-wrote the manuscript. NIN, and SR were responsible for project funding. All listed authors have read and approved the manuscript.

## COMPLIANCE WITH ETHICAL STANDARDS

The authors declare that they have no conflict of interest.

## SUPPLEMENTARY FIGURES

**Figure S1:**
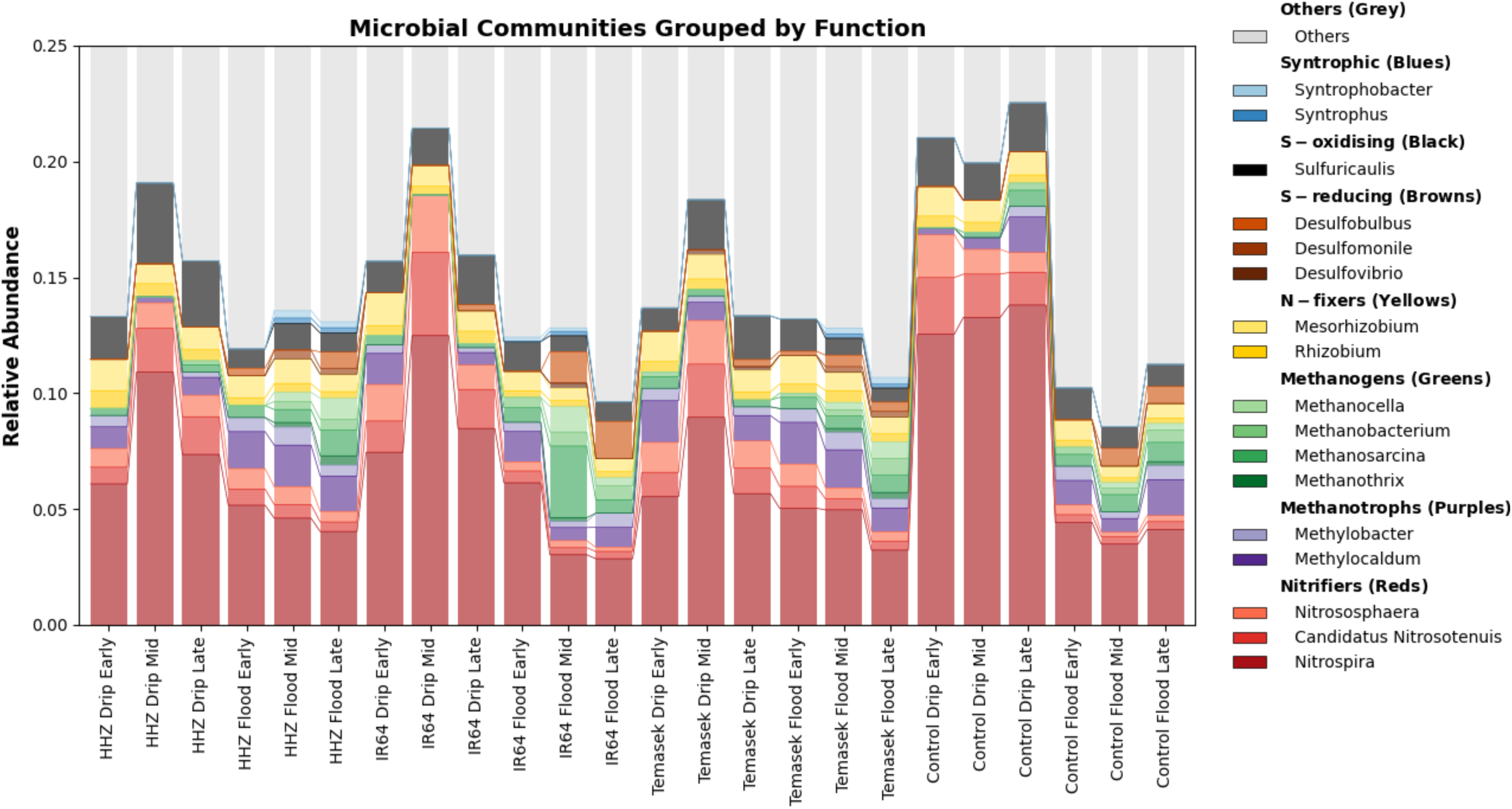
Stacked bar chart illustrating the relative abundance of methanogens, *Nitrospira* and sulphur-reducing bacteria under different irrigation methods. Methanogens and sulphur-reducing taxa like *Desulfuromonas* and *Desulfobulbus* are higher in relative abundance in flooded soil. *Nitrospira* and sulphur-oxidiser like *Sulfurcaulis* are higher in drip irrigation. *Syntrophus* and *Syntrophobacter* are also present under flooded conditions and are found in plots with rice plants.

**Figure S2:**
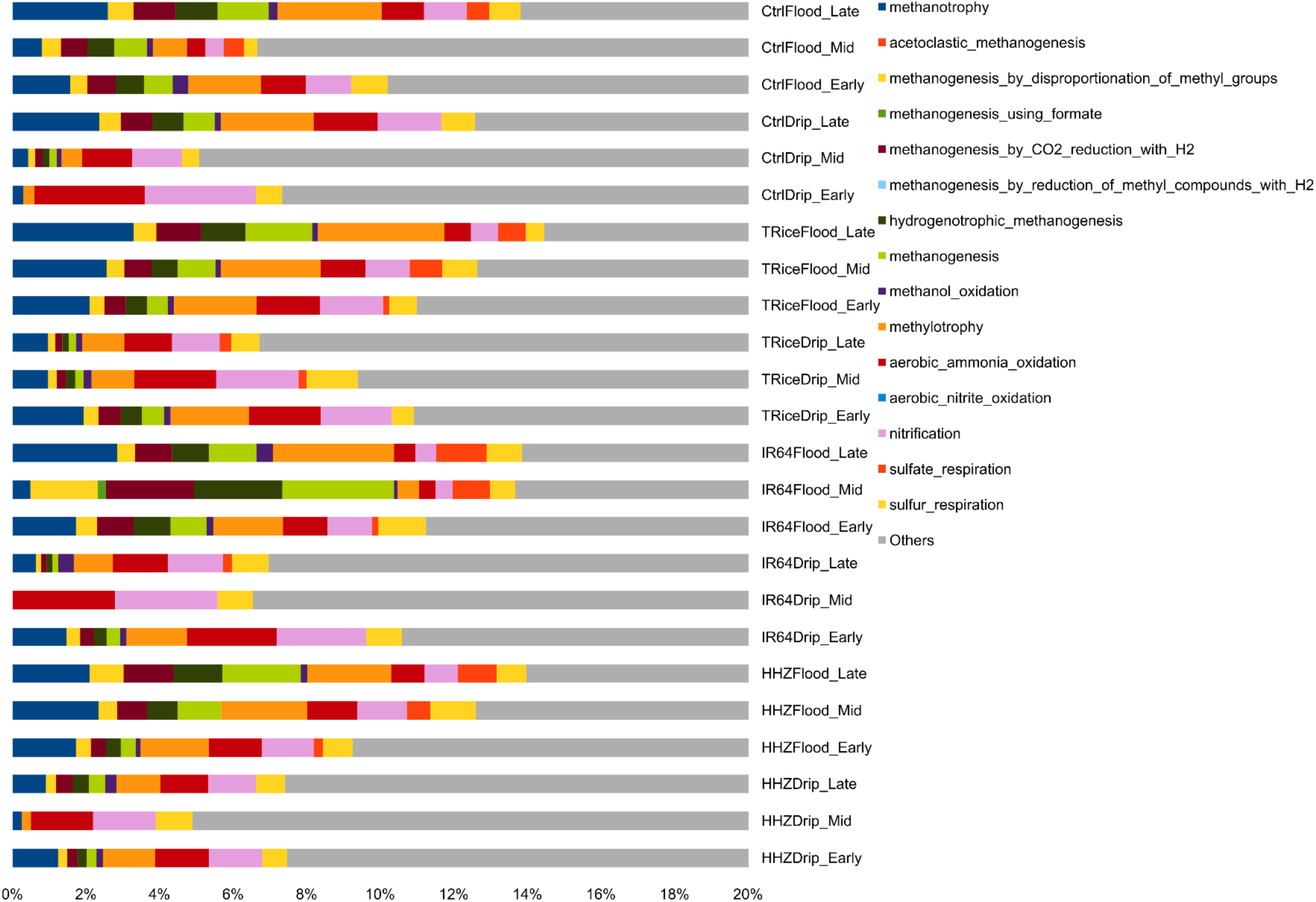
Relative abundance of predicted microbial metabolic functions across rice genotypes, irrigation treatments, and growth stages. Bars represent treatment-timepoint combinations where colours denote distinct pathways including methanogenesis, sulfur respiration, and nitrification.

**Figure S3:**
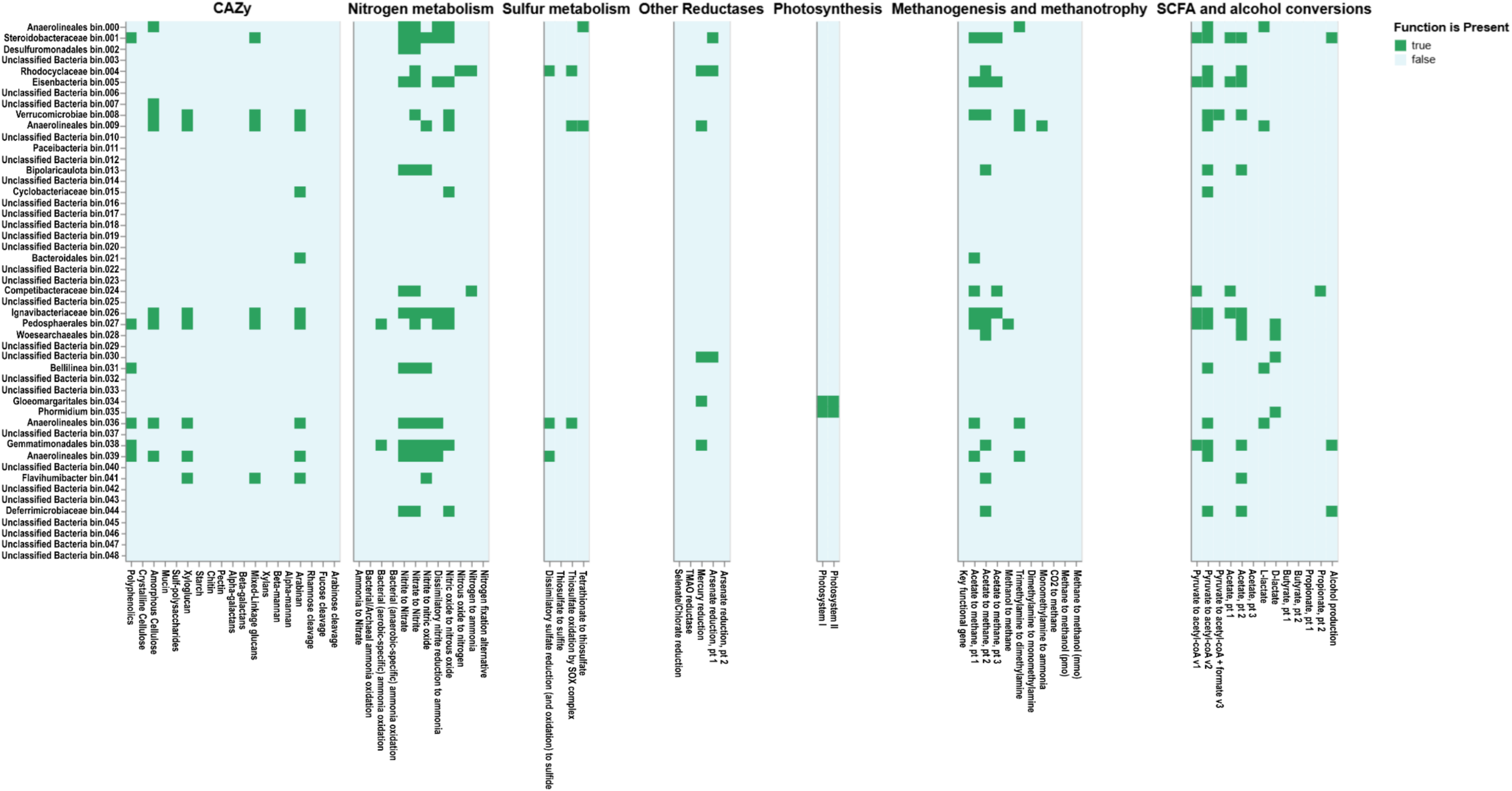
Putative functions in flood-irrigated soil. Metagenome-assembled genome bins are indicative of nitrate-to-nitrite reduction in *Rhodocyclaceae*, *Pedosphaerales* and *Gemmatimonadales*, alongside photosynthetic activity in cyanobacteria like *Phormidium* and *Gloeomargaritales* and methanogenesis-related activities in *Rhodocyclaceae* and *Pedosphaerales*.

**Figure S4:**
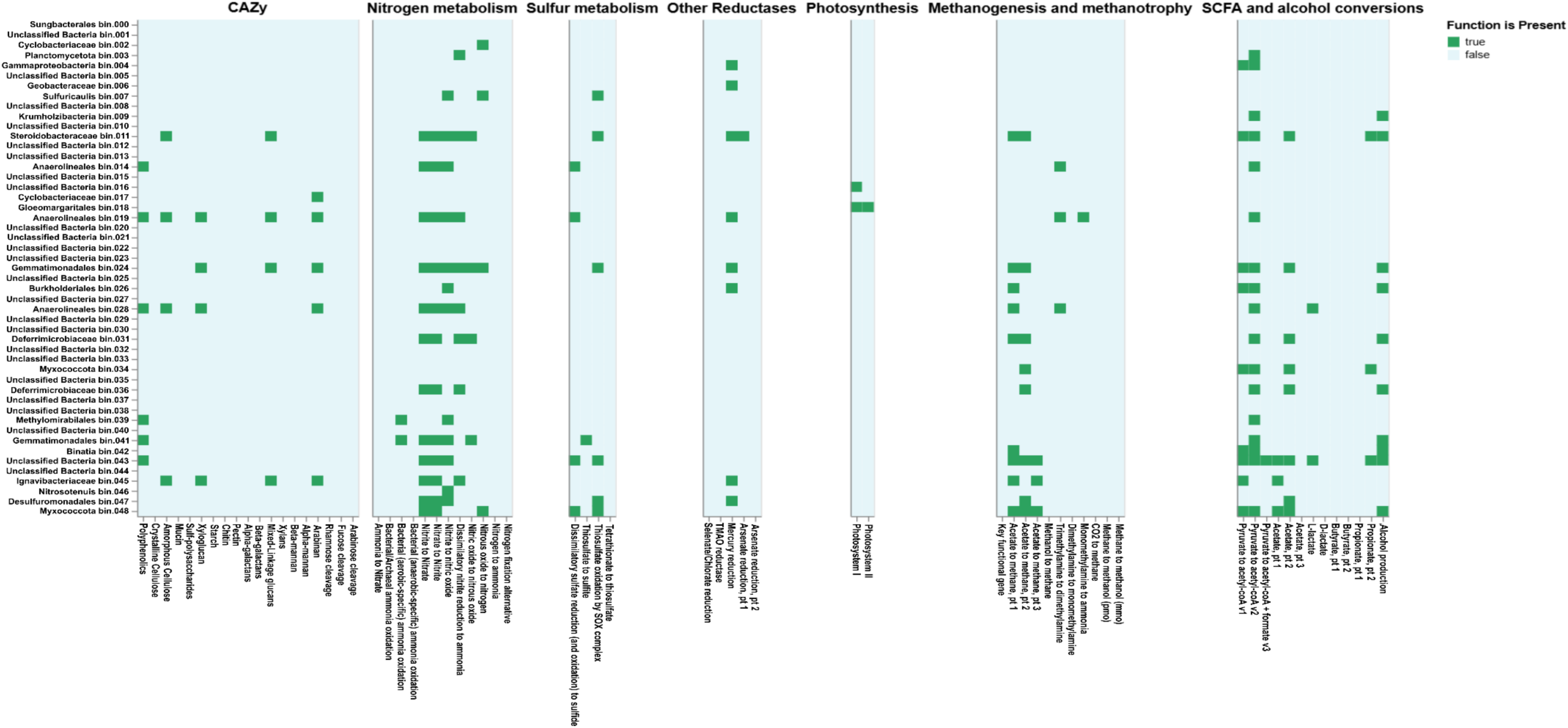
Putative functions in drip-irrigated soil. Metagenome-assembled genome bins in drip-irrigated soil highlight denitrification genes from nitrate, nitrite, nitric oxide to nitrous oxide in *Deferrimicrobiaceae* and *Gemmatimonadales*. *Nitrosotenuis* is likely involved in ammonia oxidation, *Sulfuricaulis* and *Desulfuromonadales* in sulfur cycling, while *Methylomirabilales* in methylotrophy-related activities.

**Table S1:**
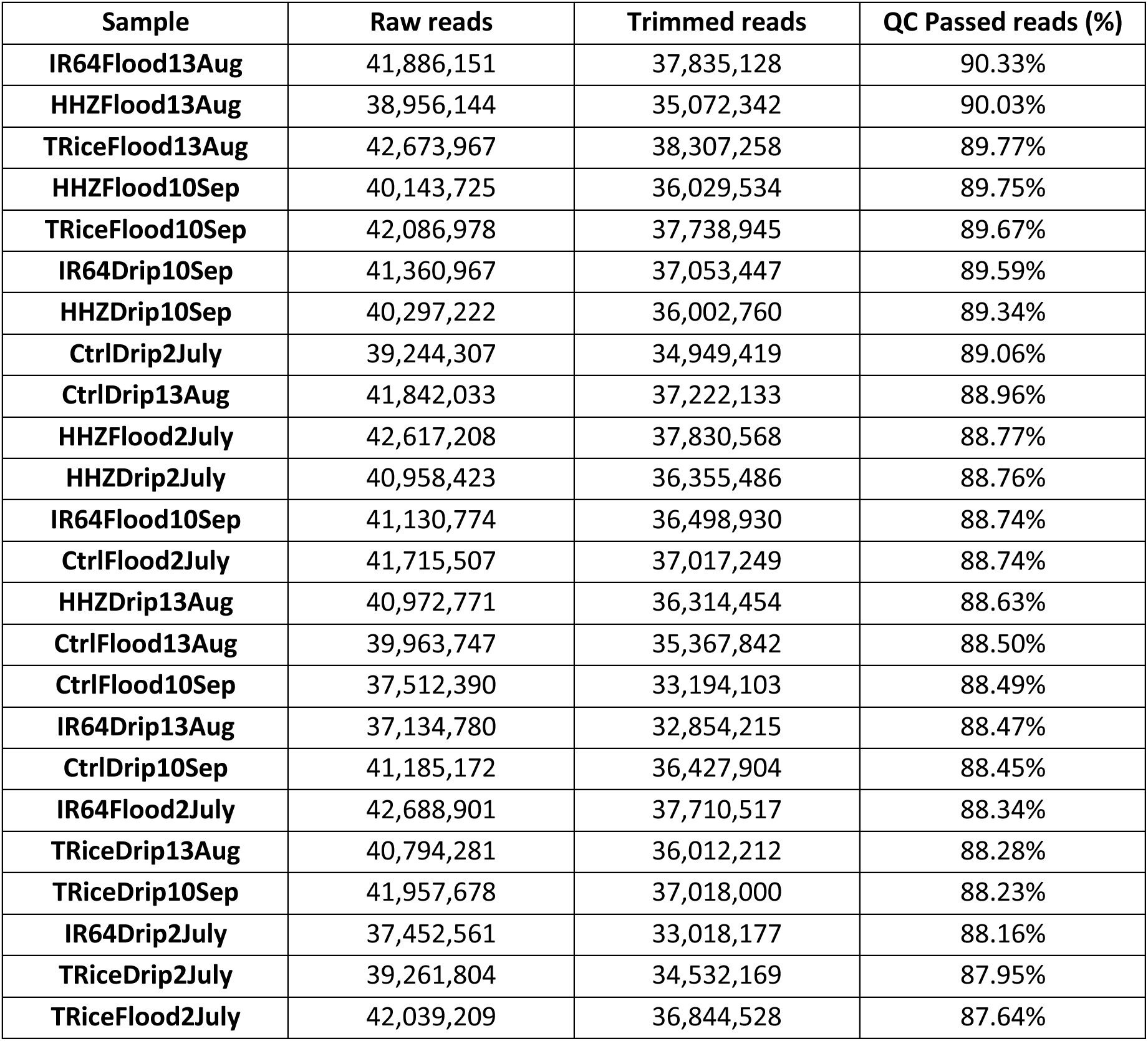
Quality of the metagenomic sequencing data from raw reads to adapter trimming and quality filtering as QC passed reads in percentage.

**Table S2:**
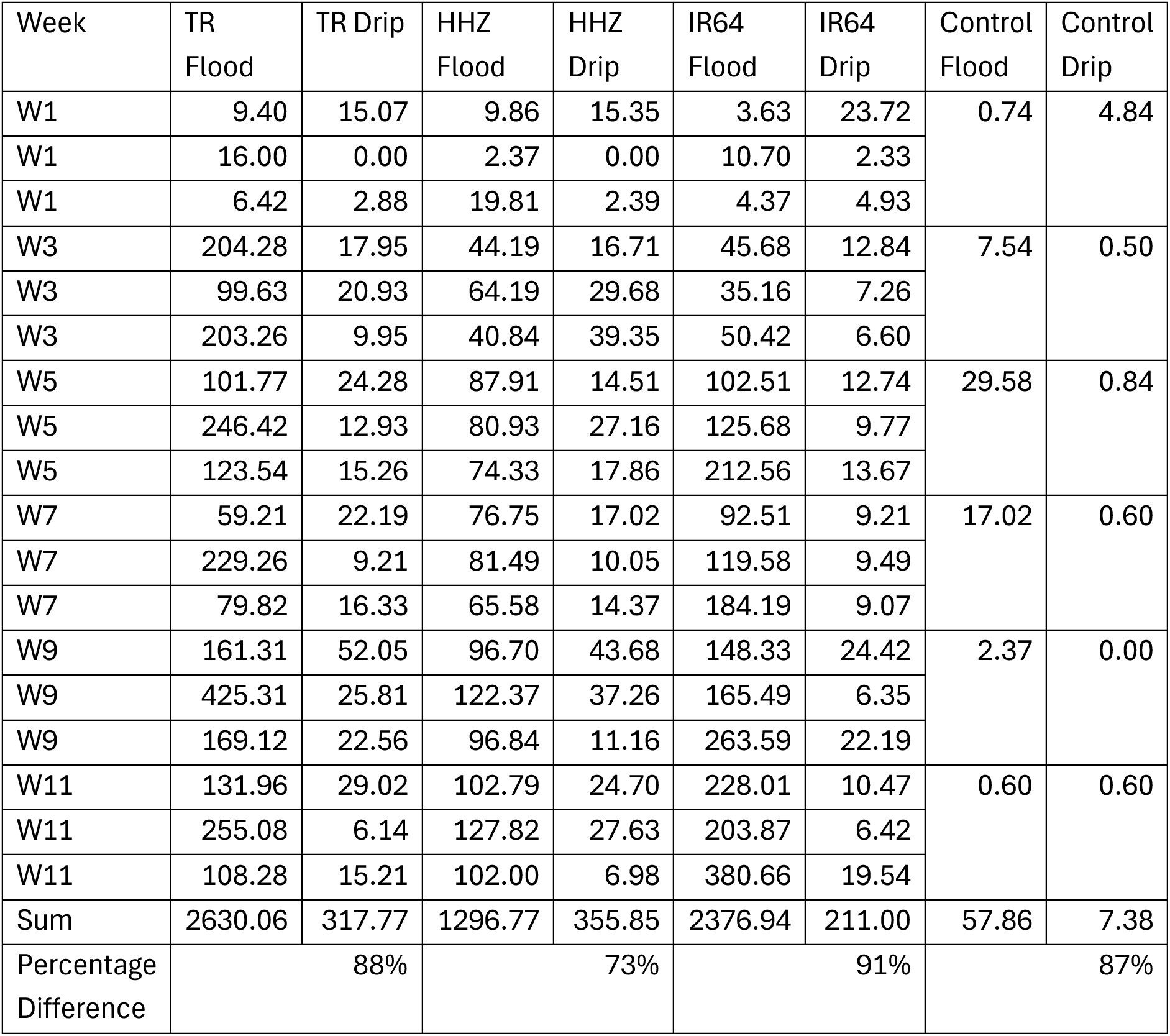
Methane emission readings from flood and drip in Temasek rice, Huanghuazhan, IR64 and blank control plots.

**Table S3:**
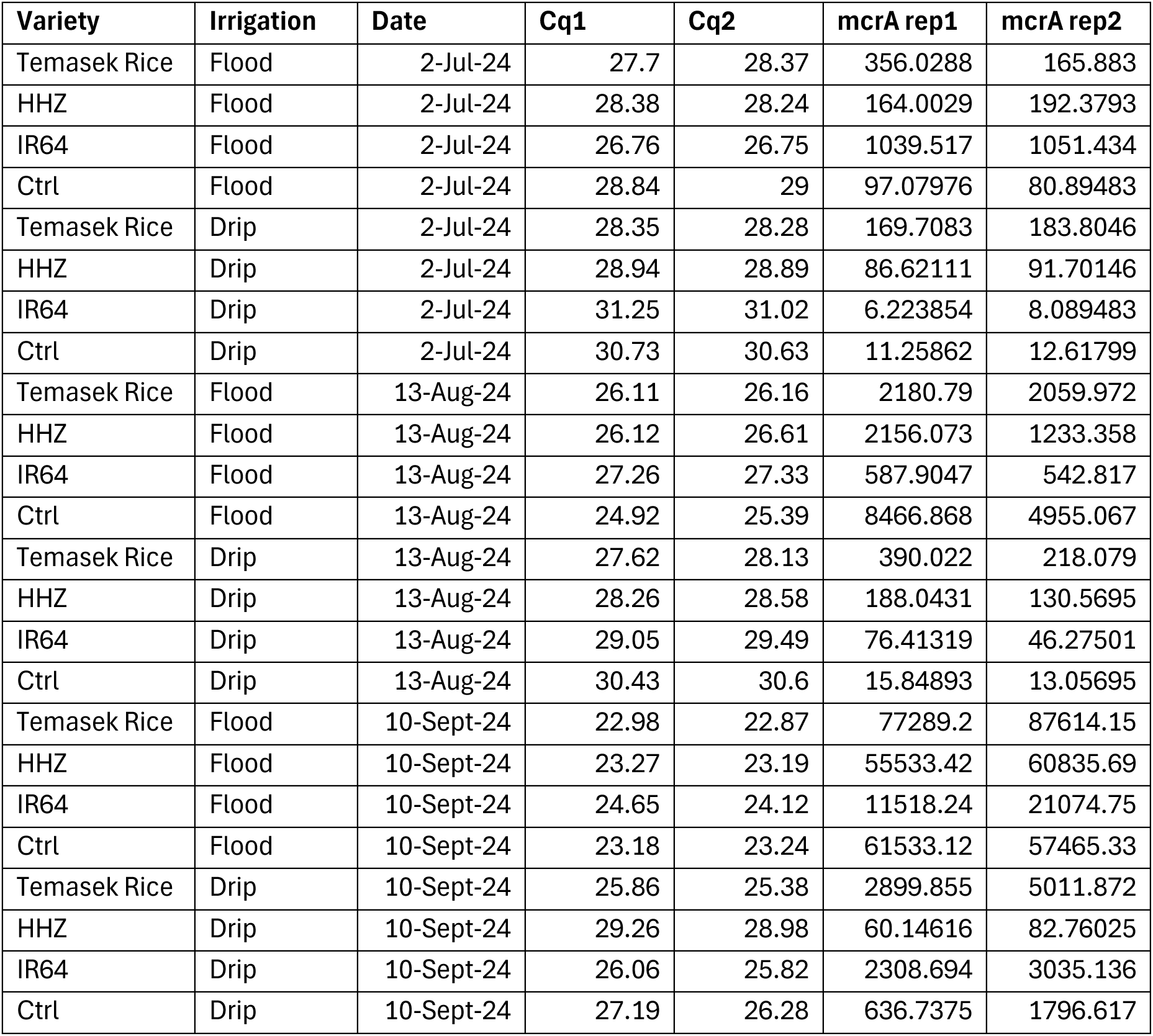
*mcrA* gene copy number estimates based on quantitative PCR assay.

**Table S4:**
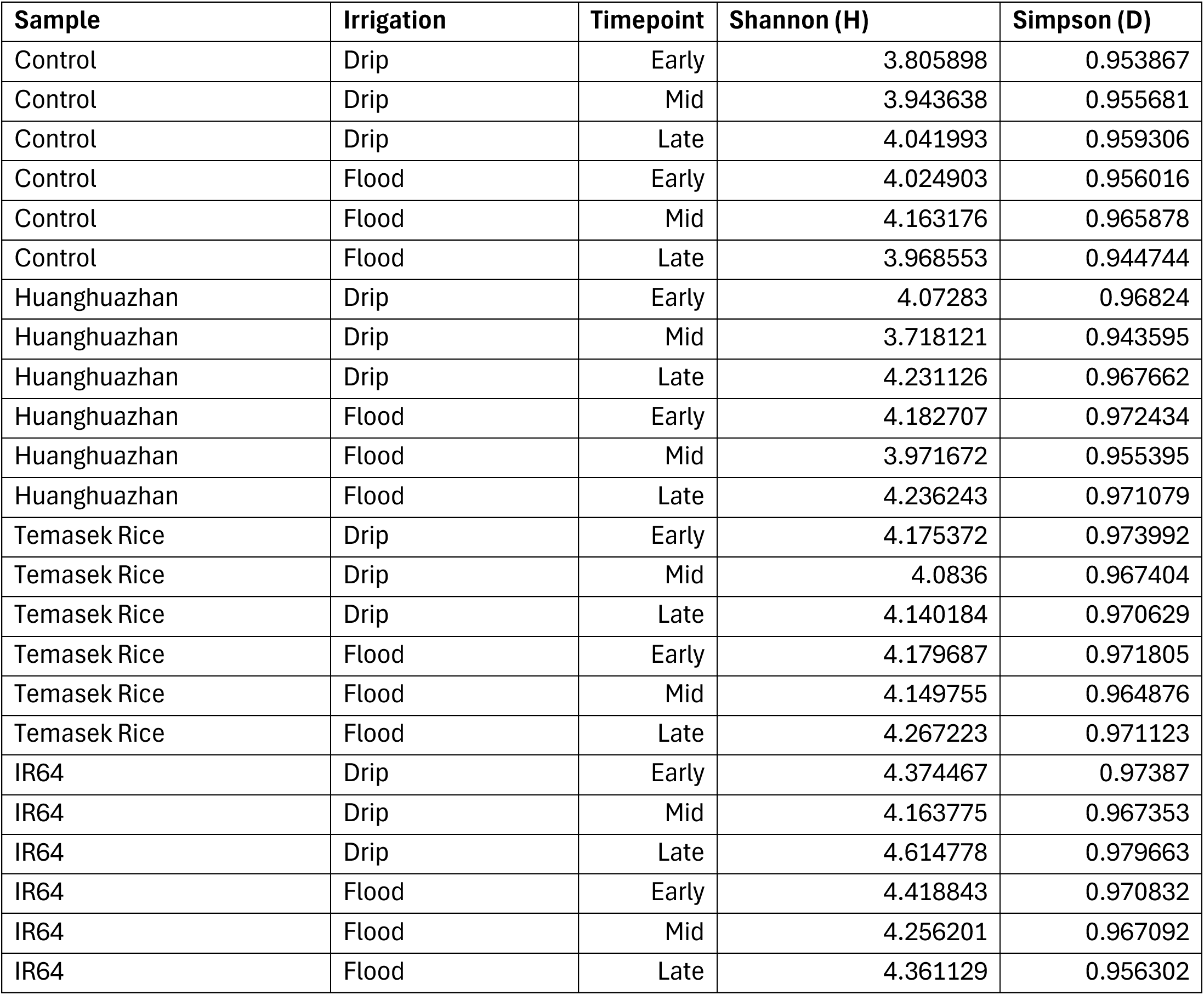
Alpha diversity of control, Huanghuazhan, Temasek Rice and IR64 under drip and flood irrigation conditions.

**Table S5:**
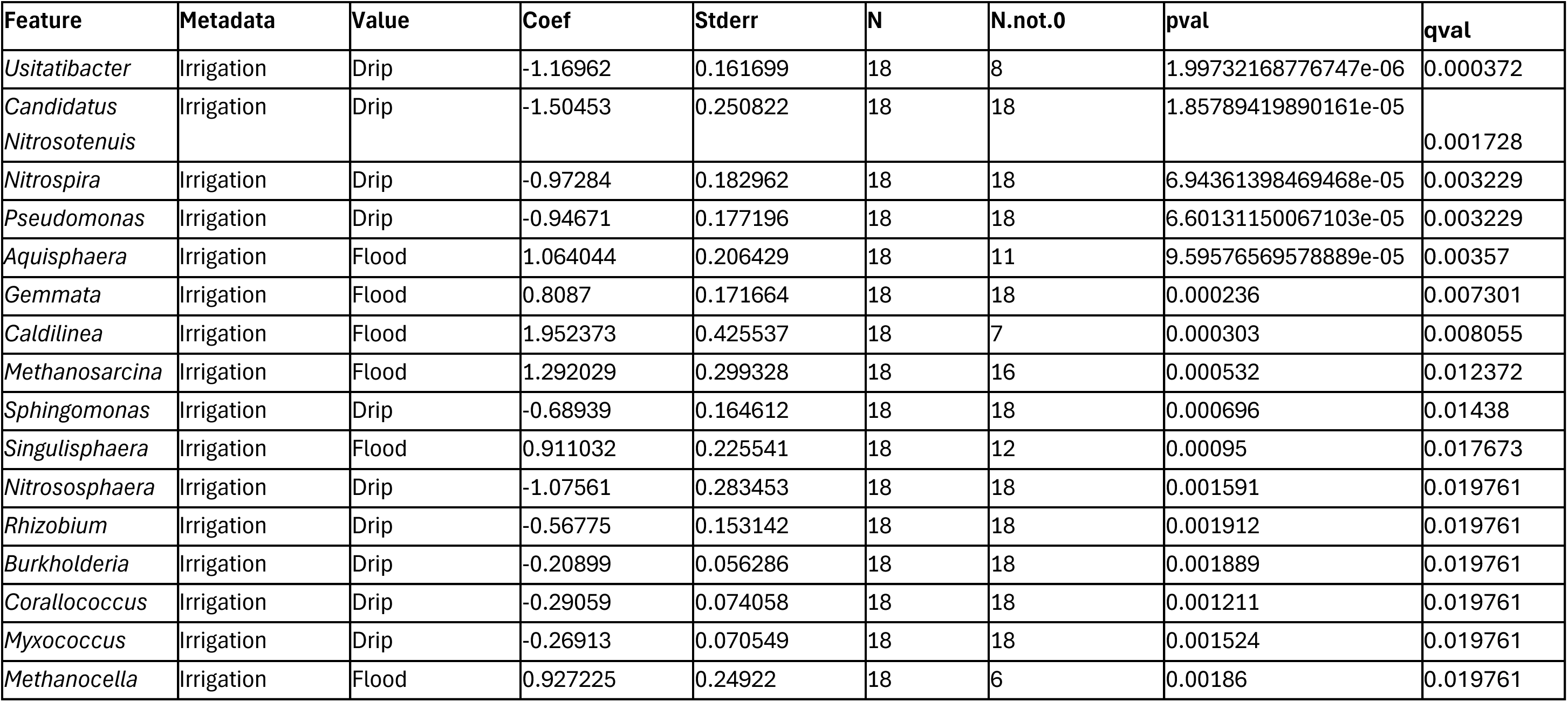
Top 16 differential taxa in drip and flood irrigated soil systems ranked by *p-*value in descending order by MaAsLin2.

**Table S6:**
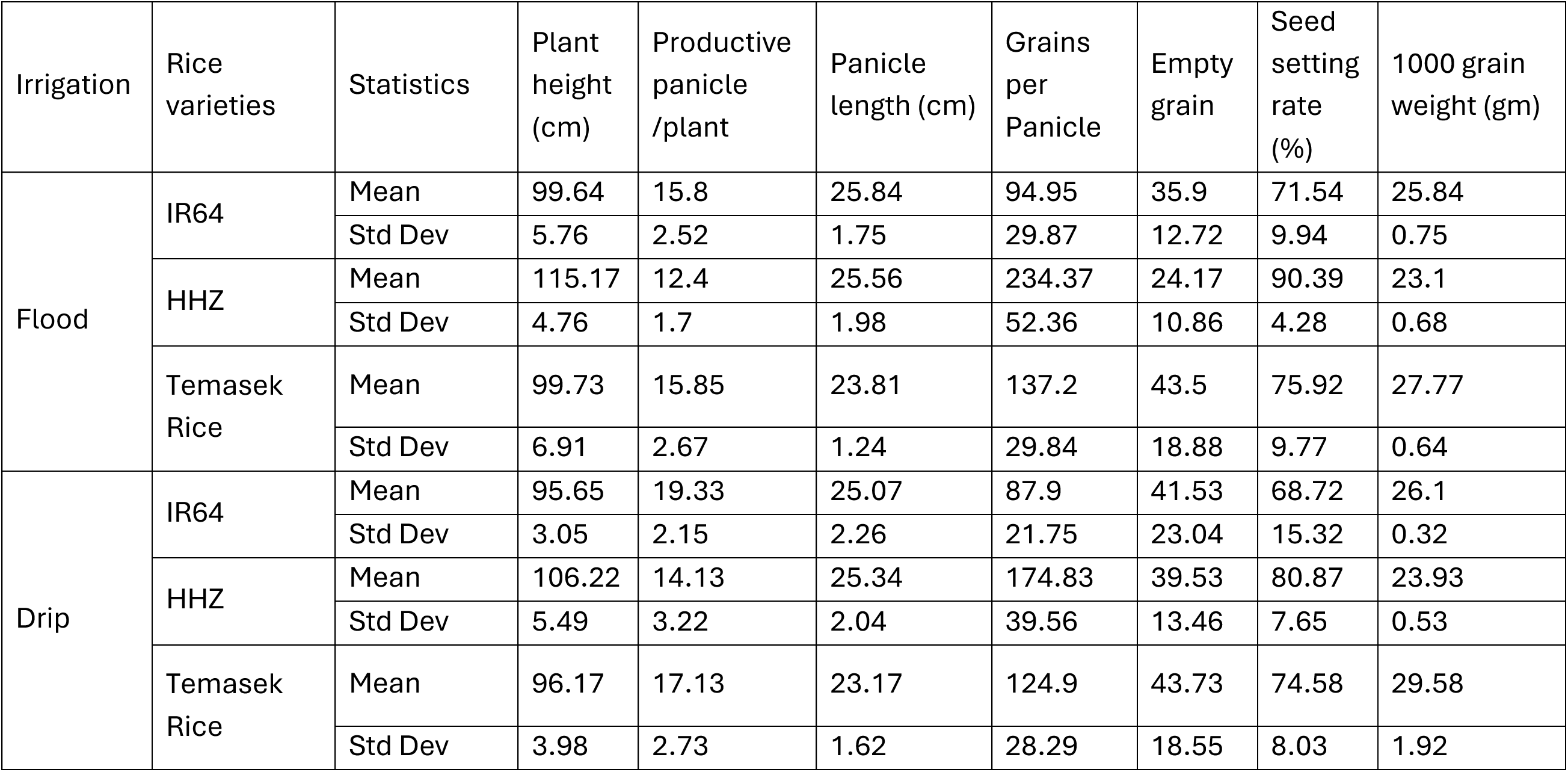
Agronomic information of rice varieties under flood and drip irrigation. Plant height, productive panicle, panicle length, grains per panicle, empty grain, seed setting rate, 1000 grain weight and yield were measured and quantified for IR64, HHZ and Temasek rice.

## REFERENCES

1. Huang J, Wang Y and Wang J. 2015. Farmers’ adaptation to extreme weather events through farm management and its impacts on the mean and risk of rice yield in China. American Journal of Agricultural Economics 97(2):602–617. 10.1093/ajae/aav005

2. Wang J, Ciais P, Smith P, et al. 2023. The role of rice cultivation in changes in atmospheric methane concentration and the Global Methane Pledge. Global Change Biology 29(10):2776–2789. 10.1111/gcb.16631

3. Neue HU, Wassmann R, Lantin RS, et al. 1996. Factors affecting methane emission from rice fields. Atmospheric Environment 30(10-11):1751–1754. 10.1016/1352-2310(95)00375-4

4. Yang Y, Shen L, Agathokleous E, et al. 2024. The interplay of soil physicochemical properties, methanogenic diversity, and abundance governs methane production potential in paddy soil subjected to multi-decadal straw incorporation. Environmental research 256:119246. 10.1016/j.envres.2024.119246

5. Kim WJ, Bui LT, Chun JB, et al. 2018. Correlation between methane (CH4) emissions and root aerenchyma of rice varieties. Plant breeding and biotechnology 6(4):381–390. 10.9787/PBB.2018.6.4.381

6. Jin Y, Liu T, Hu J, et al. 2025. Reducing methane emissions by developing low-fumarate high-ethanol eco-friendly rice. Molecular Plant 18(2):333–349. 10.1016/j.molp.2025.01.008

7. Setyanto P, Pramono A, Adriany TA, et al. 2018. Alternate wetting and drying reduces methane emission from a rice paddy in Central Java, Indonesia without yield loss. Soil Science and Plant Nutrition 64(1):23–30. 10.1080/00380768.2017.1409600

8. Tal A. 2016. Rethinking the sustainability of Israel’s irrigation practices in the Drylands. Water research 90:387–394. 10.1016/j.watres.2015.12.016

9. Conrad R. 2020. Methane production in soil environments—Anaerobic biogeochemistry and microbial life between flooding and desiccation. Microorganisms 8(6):881. 10.3390/microorganisms8060881

10. Dubey A, Malla MA, Khan F, et al. 2019. Soil microbiome: a key player for conservation of soil health under changing climate. Biodiversity and Conservation 28:2405–2429. 10.1007/s10531-019-01760-5

11. Sahu N, Vasu D, Sahu A, et al. 2017. Strength of microbes in nutrient cycling: a key to soil health. Agriculturally important microbes for sustainable agriculture: Volume I: Plant-soil-microbe nexus 69–86. 10.1007/978-981-10-5589-8_4

12. Luo Y, Zakaria S, Basyah B, et al. 2014. Marker-assisted breeding of Indonesia local rice variety Siputeh for semi-dwarf phonetype, good grain quality and disease resistance to bacterial blight. Rice 7:1–8. 10.1186/s12284-014-0033-2

13. Susanto U, Barokah U and Ali J. 2020. Correlation of agronomic and leaf traits to rice yield among Huanghuazhan variety derived lines. IOP Conference Series: Earth and Environmental Science 423(1):012034. 10.1088/1755-1315/423/1/012034

14. Mackill DJ and Khush GS. 2018. IR64: a high-quality and high-yielding mega variety. Rice 11:1–11. 10.1186/s12284-018-0208-3

15. Wickham H. ggplot2. 2011. Wiley interdisciplinary reviews: computational statistics 3(2):180–185. 10.1002/wics.147

16. Bolger AM, Lohse M and Usadel B. 2014. Trimmomatic: a flexible trimmer for Illumina sequence data. Bioinformatics 30(15):2114–2120. 10.1093/bioinformatics/btu170

17. Buchfink B, Reuter K and Drost HG. 2021. Sensitive protein alignments at tree-of-life scale using DIAMOND. Nature methods 18(4):366–368. 10.1038/s41592-021-01101-x

18. Huson DH, Auch AF, Qi J, et al. 2007. MEGAN analysis of metagenomic data. Genome research 17(3):377–386. 10.1101/gr.5969107

19. Oksanen J, Kindt R, Legendre P, et al. 2007. The vegan package. Community ecology package 10(631-637):719.

20. McMurdie PJ and Holmes S. 2013. phyloseq: an R package for reproducible interactive analysis and graphics of microbiome census data. PloS one 8(4):e61217. 10.1371/journal.pone.0061217

21. Mallick H, Rahnavard A, McIver LJ, et al. 2021. Multivariable association discovery in population-scale meta-omics studies. PLoS computational biology 17(11):e1009442. 10.1371/journal.pcbi.1009442

22. Csardi, G. and Nepusz, T. 2006. The igraph software. Complex syst, 1695, 1–9.

23. Su, G., Morris, J.H., Demchak, B. and Bader, G.D. 2014. Biological network exploration with Cytoscape 3. Current protocols in bioinformatics 47(1):8–13. 10.1002/0471250953.bi0813s47

24. Gustavsen, J.A., Pai, S., Isserlin, R., Demchak, B. and Pico, A.R. 2019. RCy3: Network biology using Cytoscape from within R. F1000Research 8:1774. 10.12688/f1000research.20887.3

25. Li, D., Liu, C.M., Luo, R., Sadakane, K. and Lam, T.W. 2015. MEGAHIT: an ultra-fast single-node solution for large and complex metagenomics assembly via succinct de Bruijn graph. Bioinformatics 31(10):1674–1676. 10.1093/bioinformatics/btv033

26. Alneberg J, Bjarnason BS, de Bruijn I, et al. 2014. Binning metagenomic contigs by coverage and composition. Nature methods 11(11):1144–1146. 10.1038/nmeth.3103

27. Parks DH, Imelfort M, Skennerton CT, et al. 2015. CheckM: assessing the quality of microbial genomes recovered from isolates, single cells, and metagenomes. Genome research 25(7):1043–1055. 10.1101/gr.186072.114

28. Chivian D, Jungbluth SP, Dehal PS, et al. 2023. Metagenome-assembled genome extraction and analysis from microbiomes using KBase. Nature Protocols 18(1):208–238. 10.1038/s41596-022-00747-x

29. Shaffer M, Borton MA, Bolduc B, et al. 2023. kb_DRAM: annotation and metabolic profiling of genomes with DRAM in KBase. Bioinformatics 39(4):btad110. 10.1093/bioinformatics/btad110

30. Ferry JG. 2020. *Methanosarcina acetivorans*: a model for mechanistic understanding of aceticlastic and reverse methanogenesis. Frontiers in Microbiology. 11:1806. 10.3389/fmicb.2020.01806

31. Jetten MS, Stams AJ and Zehnder AJ. 1992. Methanogenesis from acetate: a comparison of the acetate metabolism in *Methanothrix soehngenii* and *Methanosarcina* spp. FEMS Microbiology Reviews 8(3-4):181–197. 10.1111/j.1574-6968.1992.tb04987.x

32. Kitamura K, Fujita T, Akada S, et al. 2011. *Methanobacterium kanagiense* sp. nov., a hydrogenotrophic methanogen, isolated from rice-field soil. International journal of systematic and evolutionary microbiology 61(6):1246–1252. 10.1099/ijs.0.026013-0

33. Lü Z and Lu Y. 2012. *Methanocella conradii* sp. nov., a thermophilic, obligate hydrogenotrophic methanogen, isolated from Chinese rice field soil. PloS one 7(4):e35279. 10.1371/journal.pone.0035279

34. Liu P, Pommerenke B and Conrad R. 2018. Identification of *Syntrophobacteraceae* as major acetate-degrading sulfate reducing bacteria in Italian paddy soil. Environmental Microbiology 20(1):337–354. 10.1111/1462-2920.14001

35. Lueders T, Pommerenke B and Friedrich MW. 2004. Stable-isotope probing of microorganisms thriving at thermodynamic limits: syntrophic propionate oxidation in flooded soil. Applied and Environmental Microbiology 70(10):5778–5786. 10.1128/AEM.70.10.5778-5786.2004

36. Pan X, Li H, Zhao L, et al. 2021. Response of syntrophic bacterial and methanogenic archaeal communities in paddy soil to soil type and phenological period of rice growth. Journal of cleaner production 278:123418. 10.1016/j.jclepro.2020.123418

37. Muroski JM, Fu JY, Nguyen HH, et al. 2022. The acyl-proteome of *Syntrophus aciditrophicus* reveals metabolic relationships in benzoate degradation. Molecular & Cellular Proteomics 21(4). 10.1016/j.mcpro.2022.100215

38. Kern J, Hellebrand HJ, Gömmel M, et al. 2012. Effects of climatic factors and soil management on the methane flux in soils from annual and perennial energy crops. Biology and Fertility of Soils. 48(1):1–8. 10.1007/s00374-011-0603-z

39. Singh A and Dubey SK. 2012. Temporal variation in methanogenic community structure and methane production potential of tropical rice ecosystem. Soil Biology and Biochemistry 48:162–166. 10.1016/j.soilbio.2012.01.022

40. Ali MA, Lee CH, Lee YB, et al. 2009. Silicate fertilization in no-tillage rice farming for mitigation of methane emission and increasing rice productivity. Agriculture, ecosystems & environment 132(1-2):16–22. 10.1016/j.agee.2009.02.014

41. Lau KJX, Ma A, Chen B, et al. 2025. Controlled irrigation suppresses methane emissions by reshaping the rhizosphere microbiomes in rice. Microbiology Spectrum 02970–25. 10.1128/spectrum.02970-25

42. Chen S, Wang L, Zhang S, et al. 2023. Soil organic carbon stability mediate soil phosphorus in greenhouse vegetable soil by shifting phoD-harboring bacterial communities and keystone taxa. Science of the Total Environment 873:162400. 10.1016/j.scitotenv.2023.162400

43. Sauder LA, Engel K, Lo CC, et al. 2018. “Candidatus Nitrosotenuis aquarius,” an ammonia-oxidizing archaeon from a freshwater aquarium biofilter. Applied and Environmental Microbiology 84(19): e01430–18. 10.1128/AEM.01430-18

44. Hu J, Zhao Y, Yao X, et al. 2021. Dominance of comammox *Nitrospira* in soil nitrification. Science of the Total Environment 780:146558. 10.1016/j.scitotenv.2021.146558

45. Gamble TN, Betlach MR and Tiedje JM. 1977. Numerically dominant denitrifying bacteria from world soils. Applied and Environmental Microbiology 33(4):926–939. 10.1128/aem.33.4.926-939.1977

46. Sørensen J and Nybroe O. 2004. *Pseudomonas* in the soil environment. In Pseudomonas: Volume 1 Genomics, Life Style and Molecular Architecture 369–401. 10.1007/978-1-4419-9086-0_12

47. Zhang N, Dong C, Li L, et al. 2024. Rare bacterial and fungal taxa respond strongly to combined inorganic and organic fertilization under short-term conditions. Applied Soil Ecology 203:105639. 10.1016/j.apsoil.2024.105639

48. Singh N, Singh V, Rai SN, et al. 2022. Metagenomic analysis of garden soil-derived microbial consortia and unveiling their metabolic potential in mitigating toxic hexavalent chromium. Life 12(12):2094. 10.3390/life12122094

49. Kuo J, Liu D, Wen WH, et al. 2024. Different microbial communities in paddy soils under organic and nonorganic farming. Brazilian Journal of Microbiology 55(1):777–788. 10.1007/s42770-023-01218-5

50. Roy RK, Hosseiniyan Khatibi SM, Kim SR, et al. 2025. A robust model based on root morphological and anatomical features to distinguish high and low methane emission rice varieties through machine learning approaches. in silico Plants 7(2):diaf017. 10.1093/insilicoplants/diaf017

51. Fernandez-Baca CP, Rivers AR, Kim W, et al. 2021. Changes in rhizosphere soil microbial communities across plant developmental stages of high and low methane emitting rice genotypes. Soil Biology and Biochemistry 156:108233. 10.1016/j.soilbio.2021.108233

52. Aulakh MS, Bodenbender J, Wassmann R, et al. 2000. Methane transport capacity of rice plants. II. Variations among different rice cultivars and relationship with morphological characteristics. Nutrient Cycling in Agroecosystems 58(1):367–375. 10.1023/A:1009839929441

53. Denier van Der Gon, H.A.C., Kropff, M.J., Van Breemen, N., et al. 2002. Optimizing grain yields reduces CH4 emissions from rice paddy fields. Proceedings of the National Academy of Sciences 99(19):12021–12024. 10.1073/pnas.192276599

54. Kwon, Y., Lee, J.Y., Choi, J., et al. 2023. Loss-of-function gs3 allele decreases methane emissions and increases grain yield in rice. Nature Climate Change 13(12):1329–1333. 10.1038/s41558-023-01872-5

55. Liang B, Wang LY, Mbadinga, SM, et al. 2015. *Anaerolineaceae* and *Methanosaeta* turned to be the dominant microorganisms in alkanes-dependent methanogenic culture after long-term of incubation. Amb Express 5(1):37. 10.1186/s13568-015-0117-4

56. Ji JH, Liu YF, Zhou L, et al. 2019. Methanogenic degradation of long n-alkanes requires fumarate-dependent activation. Applied and environmental microbiology 85(16):e00985–19. 10.1128/AEM.00985-19

57. Ouboter HT, Mesman R, Sleutels T, et al. 2024. Mechanisms of extracellular electron transfer in anaerobic methanotrophic archaea. Nature Communications 15(1):1477. 10.1038/s41467-024-45758-2

58. Podosokorskaya OA, Kadnikov VV, Gavrilov SN, et al. 2013. Characterization of *Melioribacter roseus* gen. nov., sp. nov., a novel facultatively anaerobic thermophilic cellulolytic bacterium from the class *Ignavibacteria*, and a proposal of a novel bacterial phylum *Ignavibacteriae*. Environmental microbiology 15(6):1759–1771. 10.1111/1462-2920.12067

59. Zhou J, Smith JA, Li M, et al. 2023. Methane production by *Methanothrix thermoacetophila* via direct interspecies electron transfer with *Geobacter metallireducens*. MBio 14(4):e00360–23. 10.1128/mbio.00360-23

60. Singh DP, Prabha R, Yandigeri MS, et al. 2011. Cyanobacteria-mediated phenylpropanoids and phytohormones in rice (Oryza sativa) enhance plant growth and stress tolerance. Antonie Van Leeuwenhoek 100(4):557–568. 10.1007/s10482-011-9611-0

61. Liu X, Wang Y and Gu JD. 2021. Ecological distribution and potential roles of Woesearchaeota in anaerobic biogeochemical cycling unveiled by genomic analysis. Computational and Structural Biotechnology Journal 19:794–800. 10.1016/j.csbj.2021.01.013

62. Zou D, Zhang C, Liu Y, et al. 2024. Biogeographical distribution and community assembly of *Myxococcota* in mangrove sediments. Environmental Microbiome 19(1):47. 10.1186/s40793-024-00593-2

63. Rasmussen AN, Tolar BB, Bargar JR, et al. 2024. Diverse and unconventional methanogens, methanotrophs, and methylotrophs in metagenome-assembled genomes from subsurface sediments of the Slate River floodplain, Crested Butte, CO, USA. Msystems 9(7):e00314–24. 10.1128/msystems.00314-24

64. Nie WB, Xie GJ, Tan X, et al. 2023. Microbial niche differentiation during nitrite-dependent anaerobic methane oxidation. Environmental Science & Technology 57(17):7029–7040. 10.1021/acs.est.2c08094

