## Supplementary Figures & Tables for "Syntrophic microbiomes associated with methane-suppressive irrigation in rice"

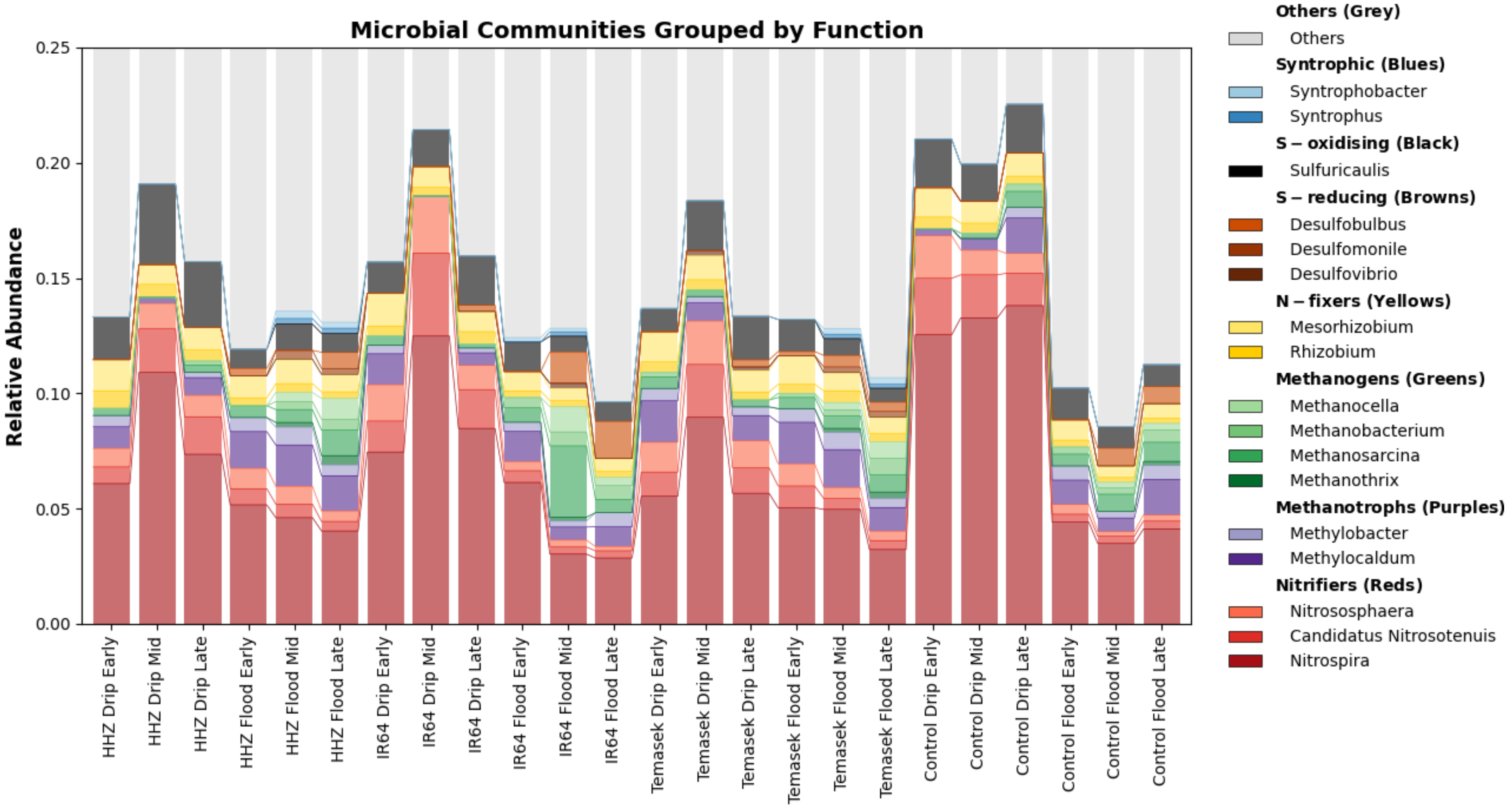


**Figure S1:** Stacked bar chart illustrating the relative abundance of methanogens, *Nitrospira* and sulphur-reducing bacteria under different irrigation methods. Methanogens and sulphur-reducing taxa like *Desulfuromonas* and *Desulfobulbus* are higher in relative abundance in flooded soil. *Nitrospira* and sulphur-oxidiser like *Sulfurcaulis* are higher in drip irrigation. *Syntrophus* and *Syntrophobacter* are also present under flooded conditions and are found in plots with rice plants.

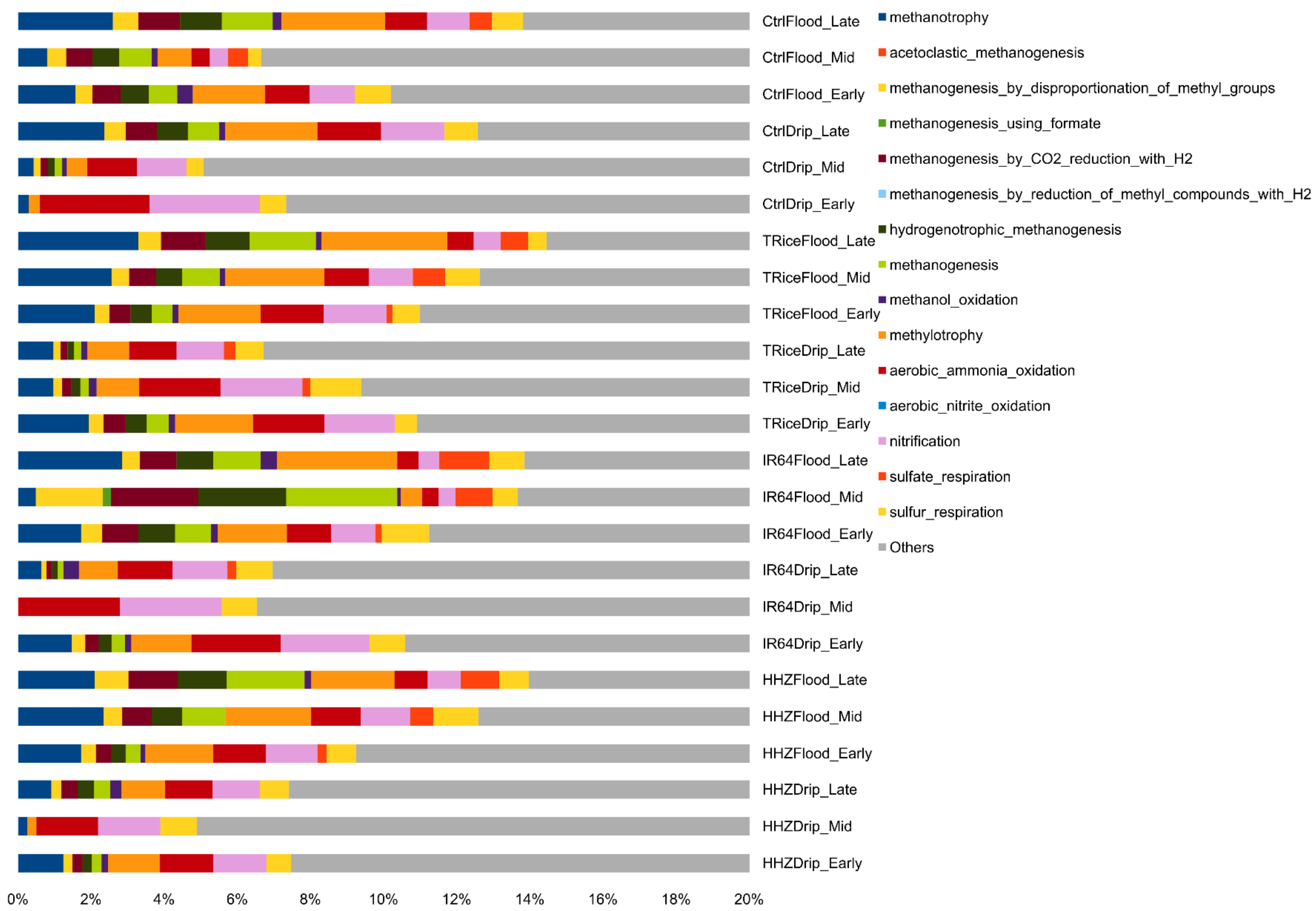


**Figure S2:** Relative abundance of predicted microbial metabolic functions across rice genotypes, irrigation treatments, and growth stages. Bars represent treatment-timepoint combinations where colours denote distinct pathways including methanogenesis, sulfur respiration, and nitrification.

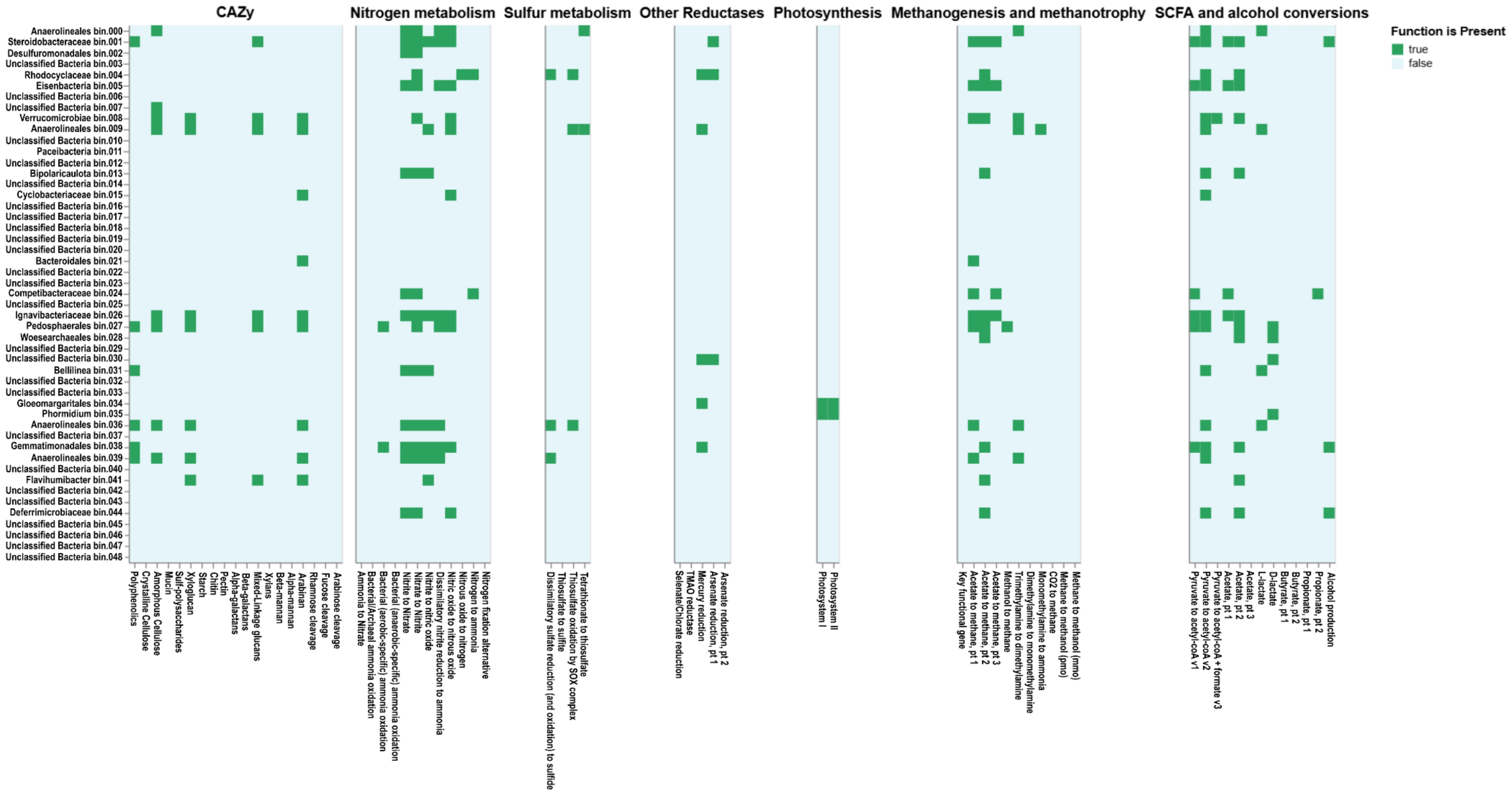


**Figure S3:** Putative functions in flood-irrigated soil. Metagenome-assembled genome bins are indicative of nitrate-to-nitrite reduction in *Rhodocyclaceae*, *Pedosphaerales* and *Gemmatimonadales*, alongside photosynthetic activity in cyanobacteria like *Phormidium* and *Gloeomargaritales* and methanogenesis-related activities in *Rhodocyclaceae* and *Pedosphaerales*.

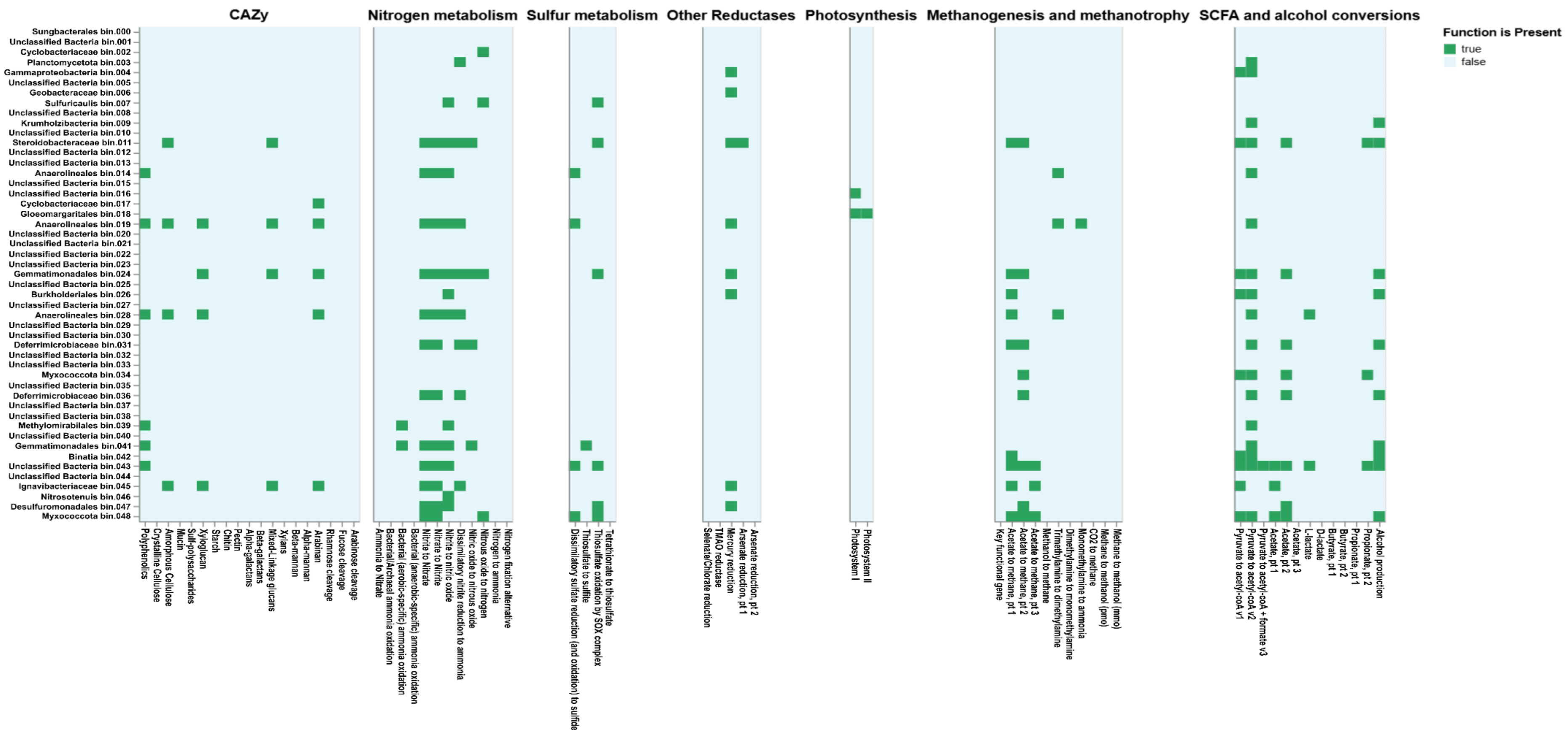


**Figure S4:** Putative functions in drip-irrigated soil. Metagenome-assembled genome bins in drip-irrigated soil highlight denitrification genes from nitrate, nitrite, nitric oxide to nitrous oxide in *Deferrimicrobiaceae* and *Gemmatimonadales*. *Nitrosotenuis* is likely involved in ammonia oxidation, *Sulfuricaulis* and *Desulfuromonadales* in sulfur cycling, while *Methylomirabilales* in methylotrophy-related activities.

**Table S1:** Quality of the metagenomic sequencing data from raw reads to adapter trimming and quality filtering as QC passed reads in percentage.

| Sample | Raw reads | Trimmed reads | QC Passed reads (%) |
|---|---|---|---|
| **IR64Flood13Aug** | 41,886,151 | 37,835,128 | 90.33% |
| **HHZFlood13Aug** | 38,956,144 | 35,072,342 | 90.03% |
| **TRiceFlood13Aug** | 42,673,967 | 38,307,258 | 89.77% |
| **HHZFlood10Sep** | 40,143,725 | 36,029,534 | 89.75% |
| **TRiceFlood10Sep** | 42,086,978 | 37,738,945 | 89.67% |
| **IR64Drip10Sep** | 41,360,967 | 37,053,447 | 89.59% |
| **HHZDrip10Sep** | 40,297,222 | 36,002,760 | 89.34% |
| **CtrlDrip2July** | 39,244,307 | 34,949,419 | 89.06% |
| **CtrlDrip13Aug** | 41,842,033 | 37,222,133 | 88.96% |
| **HHZFlood2July** | 42,617,208 | 37,830,568 | 88.77% |
| **HHZDrip2July** | 40,958,423 | 36,355,486 | 88.76% |
| **IR64Flood10Sep** | 41,130,774 | 36,498,930 | 88.74% |
| **CtrlFlood2July** | 41,715,507 | 37,017,249 | 88.74% |
| **HHZDrip13Aug** | 40,972,771 | 36,314,454 | 88.63% |
| **CtrlFlood13Aug** | 39,963,747 | 35,367,842 | 88.50% |
| **CtrlFlood10Sep** | 37,512,390 | 33,194,103 | 88.49% |
| **IR64Drip13Aug** | 37,134,780 | 32,854,215 | 88.47% |
| **CtrlDrip10Sep** | 41,185,172 | 36,427,904 | 88.45% |
| **IR64Flood2July** | 42,688,901 | 37,710,517 | 88.34% |
| **TRiceDrip13Aug** | 40,794,281 | 36,012,212 | 88.28% |
| **TRiceDrip10Sep** | 41,957,678 | 37,018,000 | 88.23% |
| **IR64Drip2July** | 37,452,561 | 33,018,177 | 88.16% |
| **TRiceDrip2July** | 39,261,804 | 34,532,169 | 87.95% |
| **TRiceFlood2July** | 42,039,209 | 36,844,528 | 87.64% |

**Table S2:** Methane emission readings from flood and drip in Temasek rice, Huanghuazhan, IR64 and blank control plots.

| Week | TR Flood | TR Drip | HHZ Flood | HHZ Drip | IR64 Flood | IR64 Drip | Control Flood | Control Drip |
|---|---|---|---|---|---|---|---|---|
| W1 | 9.40 | 15.07 | 9.86 | 15.35 | 3.63 | 23.72 | 0.74 | 4.84 |
| W1 | 16.00 | 0.00 | 2.37 | 0.00 | 10.70 | 2.33 | | |
| W1 | 6.42 | 2.88 | 19.81 | 2.39 | 4.37 | 4.93 | | |
| W3 | 204.28 | 17.95 | 44.19 | 16.71 | 45.68 | 12.84 | 7.54 | 0.50 |
| W3 | 99.63 | 20.93 | 64.19 | 29.68 | 35.16 | 7.26 | | |
| W3 | 203.26 | 9.95 | 40.84 | 39.35 | 50.42 | 6.60 | | |
| W5 | 101.77 | 24.28 | 87.91 | 14.51 | 102.51 | 12.74 | 29.58 | 0.84 |
| W5 | 246.42 | 12.93 | 80.93 | 27.16 | 125.68 | 9.77 | | |
| W5 | 123.54 | 15.26 | 74.33 | 17.86 | 212.56 | 13.67 | | |
| W7 | 59.21 | 22.19 | 76.75 | 17.02 | 92.51 | 9.21 | 17.02 | 0.60 |
| W7 | 229.26 | 9.21 | 81.49 | 10.05 | 119.58 | 9.49 | | |
| W7 | 79.82 | 16.33 | 65.58 | 14.37 | 184.19 | 9.07 | | |
| W9 | 161.31 | 52.05 | 96.70 | 43.68 | 148.33 | 24.42 | 2.37 | 0.00 |
| W9 | 425.31 | 25.81 | 122.37 | 37.26 | 165.49 | 6.35 | | |
| W9 | 169.12 | 22.56 | 96.84 | 11.16 | 263.59 | 22.19 | | |
| W11 | 131.96 | 29.02 | 102.79 | 24.70 | 228.01 | 10.47 | 0.60 | 0.60 |
| W11 | 255.08 | 6.14 | 127.82 | 27.63 | 203.87 | 6.42 | | |
| W11 | 108.28 | 15.21 | 102.00 | 6.98 | 380.66 | 19.54 | | |
| Sum | 2630.06 | 317.77 | 1296.77 | 355.85 | 2376.94 | 211.00 | 57.86 | 7.38 |
| Percentage Difference | | 88% | | 73% | | 91% | | 87% |

**Table S3:** *mcrA* gene copy number estimates based on quantitative PCR assay.

| Variety | Irrigation | Date | Cq1 | Cq2 | mcrA rep1 | mcrA rep2 |
|---|---|---|---|---|---|---|
| Temasek Rice | Flood | 2-Jul-24 | 27.7 | 28.37 | 356.0288 | 165.883 |
| HHZ | Flood | 2-Jul-24 | 28.38 | 28.24 | 164.0029 | 192.3793 |
| IR64 | Flood | 2-Jul-24 | 26.76 | 26.75 | 1039.517 | 1051.434 |
| Ctrl | Flood | 2-Jul-24 | 28.84 | 29 | 97.07976 | 80.89483 |
| Temasek Rice | Drip | 2-Jul-24 | 28.35 | 28.28 | 169.7083 | 183.8046 |
| HHZ | Drip | 2-Jul-24 | 28.94 | 28.89 | 86.62111 | 91.70146 |
| IR64 | Drip | 2-Jul-24 | 31.25 | 31.02 | 6.223854 | 8.089483 |
| Ctrl | Drip | 2-Jul-24 | 30.73 | 30.63 | 11.25862 | 12.61799 |
| Temasek Rice | Flood | 13-Aug-24 | 26.11 | 26.16 | 2180.79 | 2059.972 |
| HHZ | Flood | 13-Aug-24 | 26.12 | 26.61 | 2156.073 | 1233.358 |
| IR64 | Flood | 13-Aug-24 | 27.26 | 27.33 | 587.9047 | 542.817 |
| Ctrl | Flood | 13-Aug-24 | 24.92 | 25.39 | 8466.868 | 4955.067 |
| Temasek Rice | Drip | 13-Aug-24 | 27.62 | 28.13 | 390.022 | 218.079 |
| HHZ | Drip | 13-Aug-24 | 28.26 | 28.58 | 188.0431 | 130.5695 |
| IR64 | Drip | 13-Aug-24 | 29.05 | 29.49 | 76.41319 | 46.27501 |
| Ctrl | Drip | 13-Aug-24 | 30.43 | 30.6 | 15.84893 | 13.05695 |
| Temasek Rice | Flood | 10-Sept-24 | 22.98 | 22.87 | 77289.2 | 87614.15 |
| HHZ | Flood | 10-Sept-24 | 23.27 | 23.19 | 55533.42 | 60835.69 |
| IR64 | Flood | 10-Sept-24 | 24.65 | 24.12 | 11518.24 | 21074.75 |
| Ctrl | Flood | 10-Sept-24 | 23.18 | 23.24 | 61533.12 | 57465.33 |
| Temasek Rice | Drip | 10-Sept-24 | 25.86 | 25.38 | 2899.855 | 5011.872 |
| HHZ | Drip | 10-Sept-24 | 29.26 | 28.98 | 60.14616 | 82.76025 |
| IR64 | Drip | 10-Sept-24 | 26.06 | 25.82 | 2308.694 | 3035.136 |
| Ctrl | Drip | 10-Sept-24 | 27.19 | 26.28 | 636.7375 | 1796.617 |

**Table S4:** Alpha diversity of control, Huanghuazhan, Temasek Rice and IR64 under drip and flood irrigation conditions.

| Sample | Irrigation | Timepoint | Shannon (H) | Simpson (D) |
|---|---|---|---|---|
| Control | Drip | Early | 3.805898 | 0.953867 |
| Control | Drip | Mid | 3.943638 | 0.955681 |
| Control | Drip | Late | 4.041993 | 0.959306 |
| Control | Flood | Early | 4.024903 | 0.956016 |
| Control | Flood | Mid | 4.163176 | 0.965878 |
| Control | Flood | Late | 3.968553 | 0.944744 |
| Huanghuazhan | Drip | Early | 4.07283 | 0.96824 |
| Huanghuazhan | Drip | Mid | 3.718121 | 0.943595 |
| Huanghuazhan | Drip | Late | 4.231126 | 0.967662 |
| Huanghuazhan | Flood | Early | 4.182707 | 0.972434 |
| Huanghuazhan | Flood | Mid | 3.971672 | 0.955395 |
| Huanghuazhan | Flood | Late | 4.236243 | 0.971079 |
| Temasek Rice | Drip | Early | 4.175372 | 0.973992 |
| Temasek Rice | Drip | Mid | 4.0836 | 0.967404 |
| Temasek Rice | Drip | Late | 4.140184 | 0.970629 |
| Temasek Rice | Flood | Early | 4.179687 | 0.971805 |
| Temasek Rice | Flood | Mid | 4.149755 | 0.964876 |
| Temasek Rice | Flood | Late | 4.267223 | 0.971123 |
| IR64 | Drip | Early | 4.374467 | 0.97387 |
| IR64 | Drip | Mid | 4.163775 | 0.967353 |
| IR64 | Drip | Late | 4.614778 | 0.979663 |
| IR64 | Flood | Early | 4.418843 | 0.970832 |
| IR64 | Flood | Mid | 4.256201 | 0.967092 |
| IR64 | Flood | Late | 4.361129 | 0.956302 |

**Table S5:** Top 16 differential taxa in drip and flood irrigated soil systems ranked by *p*-value in descending order by MaAsLin2.

| Feature | Metadata | Value | Coef | Stderr | N | N.not.0 | pval | qval |
|---|---|---|---|---|---|---|---|---|
| *Usitatibacter* | Irrigation | Drip | -1.16962 | 0.161699 | 18 | 8 | 1.99732168776747e-06 | 0.000372 |
| *Candidatus Nitrosotenuis* | Irrigation | Drip | -1.50453 | 0.250822 | 18 | 18 | 1.85789419890161e-05 | 0.001728 |
| *Nitrospira* | Irrigation | Drip | -0.97284 | 0.182962 | 18 | 18 | 6.94361398469468e-05 | 0.003229 |
| *Pseudomonas* | Irrigation | Drip | -0.94671 | 0.177196 | 18 | 18 | 6.60131150067103e-05 | 0.003229 |
| *Aquisphaera* | Irrigation | Flood | 1.064044 | 0.206429 | 18 | 11 | 9.59576569578889e-05 | 0.00357 |
| *Gemmata* | Irrigation | Flood | 0.8087 | 0.171664 | 18 | 18 | 0.000236 | 0.007301 |
| *Caldilinea* | Irrigation | Flood | 1.952373 | 0.425537 | 18 | 7 | 0.000303 | 0.008055 |
| *Methanosarcina* | Irrigation | Flood | 1.292029 | 0.299328 | 18 | 16 | 0.000532 | 0.012372 |
| *Sphingomonas* | Irrigation | Drip | -0.68939 | 0.164612 | 18 | 18 | 0.000696 | 0.01438 |
| *Singulisphaera* | Irrigation | Flood | 0.911032 | 0.225541 | 18 | 12 | 0.00095 | 0.017673 |
| *Nitrososphaera* | Irrigation | Drip | -1.07561 | 0.283453 | 18 | 18 | 0.001591 | 0.019761 |
| *Rhizobium* | Irrigation | Drip | -0.56775 | 0.153142 | 18 | 18 | 0.001912 | 0.019761 |
| *Burkholderia* | Irrigation | Drip | -0.20899 | 0.056286 | 18 | 18 | 0.001889 | 0.019761 |
| *Corallococcus* | Irrigation | Drip | -0.29059 | 0.074058 | 18 | 18 | 0.001211 | 0.019761 |
| *Myxococcus* | Irrigation | Drip | -0.26913 | 0.070549 | 18 | 18 | 0.001524 | 0.019761 |
| *Methanocella* | Irrigation | Flood | 0.927225 | 0.24922 | 18 | 6 | 0.00186 | 0.019761 |

**Table S6:** Agronomic information of rice varieties under flood and drip irrigation. Plant height, productive panicle, panicle length, grains per panicle, empty grain, seed setting rate, 1000 grain weight and yield were measured and quantified for IR64, HHZ and Temasek rice.

| Irrigation | Rice varieties | Statistics | Plant height (cm) | Productive panicle /plant | Panicle length (cm) | Grains per Panicle | Empty grain | Seed setting rate (%) | 1000 grain weight (gm) |
|---|---|---|---|---|---|---|---|---|---|
| Flood | IR64 | Mean | 99.64 | 15.8 | 25.84 | 94.95 | 35.9 | 71.54 | 25.84 |
| | | Std Dev | 5.76 | 2.52 | 1.75 | 29.87 | 12.72 | 9.94 | 0.75 |
| | HHZ | Mean | 115.17 | 12.4 | 25.56 | 234.37 | 24.17 | 90.39 | 23.1 |
| | | Std Dev | 4.76 | 1.7 | 1.98 | 52.36 | 10.86 | 4.28 | 0.68 |
| | Temasek Rice | Mean | 99.73 | 15.85 | 23.81 | 137.2 | 43.5 | 75.92 | 27.77 |
| | | Std Dev | 6.91 | 2.67 | 1.24 | 29.84 | 18.88 | 9.77 | 0.64 |
| Drip | IR64 | Mean | 95.65 | 19.33 | 25.07 | 87.9 | 41.53 | 68.72 | 26.1 |
| | | Std Dev | 3.05 | 2.15 | 2.26 | 21.75 | 23.04 | 15.32 | 0.32 |
| | HHZ | Mean | 106.22 | 14.13 | 25.34 | 174.83 | 39.53 | 80.87 | 23.93 |
| | | Std Dev | 5.49 | 3.22 | 2.04 | 39.56 | 13.46 | 7.65 | 0.53 |
| | Temasek Rice | Mean | 96.17 | 17.13 | 23.17 | 124.9 | 43.73 | 74.58 | 29.58 |
| | | Std Dev | 3.98 | 2.73 | 1.62 | 28.29 | 18.55 | 8.03 | 1.92 |
